# A specific mesh-like organization of human septin octameric complex drives membrane reshaping and curvature sensitivity

**DOI:** 10.1101/2022.11.02.514824

**Authors:** Koyomi Nakazawa, Gaurav Kumar, Brieuc Chauvin, Aurélie Di Cicco, Luca Pellegrino, Michael Trichet, Bassam Hajj, João Cabral, Anirban Sain, Stéphanie Mangenot, Aurélie Bertin

## Abstract

Septins are cytoskeletal proteins interacting with the inner plasma membrane and other cytoskeletal partners. Being key in membrane remodeling processes, they often localize at specific micrometric curvatures. To analyze the behavior of human septins at the membrane, we have used a combination of methods to assay their ultrastructural organization, their curvature sensitivity as well as their role in membrane reshaping. In contrast to budding yeast septins, on membranes, human septins systematically organize into a two-layered mesh of orthogonal filaments instead of generating parallel sheets of filaments observed for budding yeast septins. This peculiar mesh organization is curvature sensitive and drives membrane reshaping as well. The observed membrane deformations together with the filamentous organization are recapitulated in a coarsegrained computed simulation to understand their mechanisms. Our results highlight the specificity of animal septins as opposed to fungal proteins.

## Introduction

Septins are ubiquitous cytoskeletal protein complexes^1^ which interact directly with the inner plasma membranes^2–4^, as well as with actin and with microtubule^5^ in eukaryotes. Septins are involved in essential cellular functions: cell division^6–8^, cell motility^9^, cell compartmentalization^10–12^. Being multi-tasking, the malfunction of septins is the cause of several diseases^13^. Septins mis-regulation are thereby associated with the emergence of cancers and neurodegenerative disorders among other major illnesses^14^. So far, the molecular organization and resulting mechanisms associated with the observed multitasking functions of septins remain essentially unknown.

In humans, 13 septin subunits are expressed in a tissue dependent manner^15^. They can assemble into linear palindromic complexes either hexameric or octameric^16^. Hence, multiple septin subunit combinations can possibly be observed. The behavior of human septins appears universal, particularly on membranes. Indeed, human septins complexes are often interacting with membranes displaying micrometer scale membrane curvatures^17^. They are found, assembling into ring-like structures, at the intercellular bridge connecting two dividing cells^18,19^, at the base of cilia^11^, at the annulus of spermatozoa^20^ or at the base of dendrites^21^. When bound to membranes, mammalian septins seem to control both the integrity and mechanical stability of membranes^9,22^. Septins are thus likely involved in membrane remodeling.

Indeed, from our previous investigations focused on budding yeast septins^23^, we have shown that they can deform membranes in a curvature sensitive manner by directly interacting with membranes. In budding yeast, septins were shown to be essential for cell division and localized at the constriction sites between the mother cell and the bud^8^. As opposed to yeast septins which have been already thoroughly studied using bottom-up in vitro assays, the behavior of human septins at the membrane has not been examined in controlled environments in vitro and barely examined in cellular contexts^16^. Indeed, until very recently human septin octamers including the essential Sept9 were not available for in vitro studies. A single study from 2006^24^ using hexameric septins (deprived of Sept9) had shown that septin complexes could produce tubulation at the surface of GUVs, suggesting that human septins could remodel membranes. Structures from individual human septin subunits had also been generated^25^. Recently, we have expressed and purified SEPT2, SEPT6, SEPT7 and SEPT9i1 complexes recombinantly from E.coli^26^.

In the present work, we have studied how human septin complexes can possibly remodel membranes. We have used a combination of bottom-up in vitro assays at different scales using fluorescence and electron microscopies. Hence, the structural organization of septins filaments coupled with membrane remodeling could be probed, simultaneously. In the present work, we found that similarly, to budding yeast septins, human septin octameric complexes are curvature sensitive and remodel membranes. However, the nature of the observed deformations as well as the human septin organization contrast with the observations gathered using budding yeast septins. We show that human septin filaments indeed assemble into a network-like array interconnecting two layers of septin filaments and the observed deformations display a curvature opposite to the ones induced by yeast septins. This distinct behavior reflects a peculiar and unique behavior of human septin complexes when bound to membranes. The observed membrane remodeling and filamentous organization was modeled by a coarse-grained simulation based on a nematic liquid crystalline 2D ordering of the septin filaments. The model confirms that this specific behavior results from the mesh size organization of human septins. Our observations thus suggest that the function of metazoan septins have significantly diverged from the role of fungi septins at the membrane.

## Results

### Self-assembly of human septins filaments on lipid model membranes

To characterize and visualize the polymerization and self-assembly of septins interacting with lipid model membranes, human septins octamers (SEPT2-GFP-SEPT6-SEPT7-SEPT9_i1-SEPT9_i1-SEPT7-SEPT6-SEPT2-GFP) were incubated with model lipid membranes. We first investigated whether negatively charged DOPS and PI(4,5)P_2_ could play a crucial role in the recruitment of septins to membranes using fluorescence microscopy. Giant unilamellar vesicles (GUVs) of different lipid compositions were generated and incubated with septins (see Method section). Septins exclusively interacted with membranes containing negatively charged lipids (DOPS (Supplementary Figure 1.B) and/or PI(4,5)P_2_ (Supplementary Figure 1)). Besides, the interaction was enhanced in the presence of both DOPS and PI(4,5)P_2_ (Supplementary Figure 1.C), which resulted in a enhanced recruitment of septins at the surface of GUVs. Interestingly, to detect any interaction between membranes and budding yeast septins, PI(4,5)P_2_ was required and DOPS was not sufficient^27^. In the following assays, artificial lipidic systems (SUV, LUV or GUVs) thus contain both DOPS and PI(4,5)P_2_. The ionic strength was adjusted to induce septins polymerization, with 75 mM NaCl. Polymerization and further filaments higher order assemblies occurred within minutes.

To visualize both vesicles and septin filaments in three dimensions with nanometer resolutions, we performed cryo-electron tomography. We observed the organization of septins bound to large unilamellar vesicles (LUVs) which are small enough to be observed by transmission cryo-electron microscopy. Indeed, GUVs, with typical micrometer size, are too thick to be suitable for observation by transmission cryo-EM. Figure 1 displays slices of cryo-electron tomograms of representative LUVs decorated with septins filaments. In most cases, orthogonal mesh-like arrangements of septin filaments were observed, bound to membranes as highlighted by the segmentations (Figures 1.A, 1B, 1.C, right panels). Interestingly, from the tomograms shown in Figure 1 and movie S1, those networks combine two layers of septins orthogonal to one another (the first and second layers are shown in blue and pink, respectively in Figure 1 bottom right). The first layer of septin filaments interacts directly with the membrane, while the second layer seems to interact with the first layer of septins. Within the mesh, the distance between filaments was measured at 18 ± 15 nm (mean ± SD, N = 157 (see histogram in Supplementary Figure 2)). Occasionally, membrane deformations are observed at nanometer scales (Figure 1 top). It is however not clear whether these nanometric deformations are caused by septins-membranes interactions. Indeed, perfectly spherical vesicles coated with septins meshes (Figure 1, middle) are also observed.

**Figure 1:**
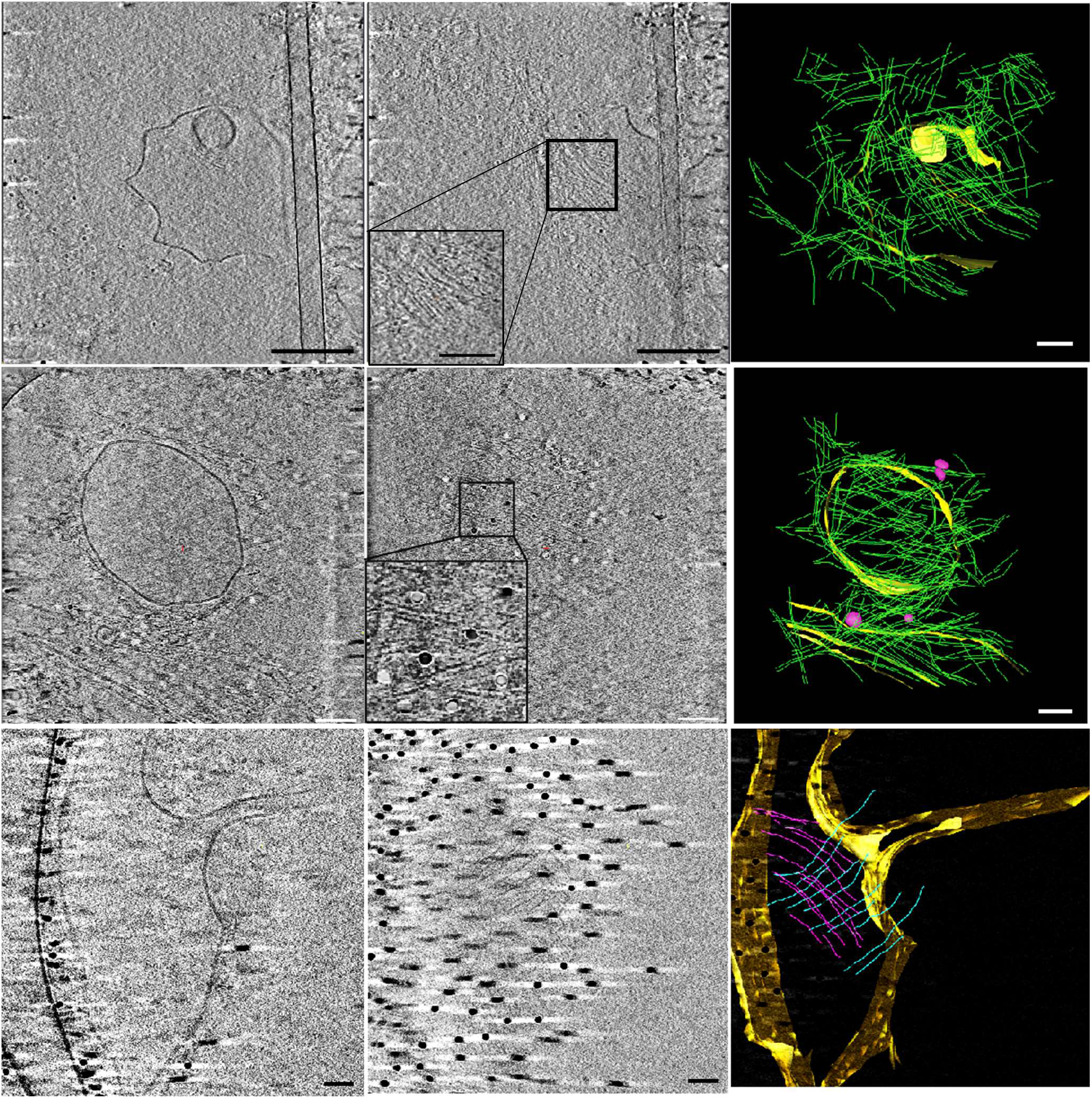
Cryo-electron tomography of septins bound to liposome. Slices of cryo-electron tomography (left and middle) and models (right) build by segmentation of tomograms. Left column displays slice where the membrane is visible. Middle column displays slice where one layer of the mesh is visualized. Septins bound to deformed LUV (top) and round shape LUV (middle) are organized as mesh structures. Mesh is constituted with two layers of septins (bottom), 1^st^ layer (bound to membrane) and 2^nd^ layer (bound to 1^st^ layer of septins filaments) of septins filaments are segmented as blue and pink color in the right image. Scale bars= 100 nm except 200 nm for top left images.

### Network of septins filaments on micrometric undulated membrane on solid surface

Budding yeast septins are known to be sensitive to micrometer curvatures^23,28^. We have thus tested whether human septins could sense curvature at micrometer scale. To this end, septin filaments were incubated with supported lipid bilayers, deposited on undulated wavy solid substrates designed to display a micrometer periodicity. Both the amplitude and the periodicity of the substrates can be finely tuned^29,30^. The resulting samples were imaged by scanning electron microscopy (SEM). Lipid bilayers strongly interact with the substrates and are thus not deformable. Our observations are presented in Figure 2. The wavy substrates displayed a periodicity of 1.6 μm and an amplitude (from peak to peak) of 200 nm, corresponding to curvatures ranging from −3 to +3 μm^-1 29^ (see Methods). These parameters were set after previous reports suggested a micrometer curvature sensitivity of octameric budding yeast septins^23,28^. Solutions of recombinant human septins octamers arranged as: SEPT2-GFP-SEPT6-SEPT7-SEPT9_i1-SEPT9_i1-SEPT7-SEPT6-SEPT2-GFP (concentrations ranging from about 10 to 100 nM) were incubated with the substrates covered with a lipid bilayer, in a 75 mM NaCl, 10 mM Tris pH 7.8 buffer. At low protein concentrations (8 nM), barely any septin filaments were detected within the convex sections of the substrates while polymerized septin filaments were observed at concave sections (Figure 2.A). Single septin filaments thereby presented a higher affinity for concave (negative) curvatures than for convex (positive) curvatures. Interestingly, the orientation of septins filaments in the concave region was specific. Indeed, septin filaments were oriented perpendicular to the longitudinal axis of 1D periodic substrate waves. At 26 nM septin concentrations (Figure 2.B), septin filaments were still present at the concave negative curvature of the substrate. Additionally, they are distributed at convex positive curvatures as well. The orientation of the filaments in the convex (positive curvature) sections was orthogonal to the orientation of the filaments present on the concave (negative curvature) sections. Hence, on convex regions, the filaments remained straight and unbent, following the direction of the periodic wavy lines of the substrate. (Figure 2.B, middle). In addition, in the same concentration range (26 nM bulk septin concentrations), local higher septin densities displaying an alternative organization were observed. This observation thus suggest that 26 nM was close to a threshold bulk septin concentration where a drastic septins re-organization was visualized (Figure 2.C and Supplementary Table 1). Indeed, septin filaments from the concave domains extended towards convex domains and were superimposed orthogonally to the straight septins filaments running along the convex hills, forming a mesh-like structures over two layers (Figure 2.C). This mesh-like organization reflected similar conclusions obtained from cryo-electron microscopy (see Figure 1). In this mesh-like arrangement, the first layer of septin filaments thus interacted directly with the membrane, while the second layer was most probably connected to the first layer of septins. Distances between adjacent septin filaments (*d*_1_ = 46 ± 20 nm and *d*_2_ = 55 ± 24 nm, Figure 2e), in the mesh-like organization, were distributed at approximatively 50 nm. In addition to this specific mesh organization, another filamentous pattern was detected. Filaments from concave domains were merged with and seen running alongside adjacent filaments on convex domains by assembling into semicircular patterns (diameters of about 0.8 μm) (Figure 1.C and Supplementary Figure 3). Above 44 nM septin concentrations (Supplementary Figure 5), both convex and concave regions of membranes were already fully covered with a meshed network of septins. To fully ensure full saturation of the surface with septins, we used a higher concentration of 90 nM (Figure 3.D). Interestingly, we could not detect any intermediate between the sparse filamentous decoration observed at 26 nM and the maximal saturation with proteins. At full saturation, the mesh-like organization in two layers was ubiquitous and thus present at both concave and convex domains (negative and positive curvature respectively, (Figure 2.D)), resulting in more regular periodicities in distances between filaments (*d*_1_ = 27 ± 11 (nm) and *d*_2_ = 19 ± 8 (nm) shown in Figure 3.E (see signals around 1/30 nm^-1^ in 2D Fourier transformation in Supplementary Figure 4). The typical spacing recovered from those dense meshes, on average, reflected the octameric periodicities of human septins (32 nm). Besides, we never observed any additional third protein layer on top of the described orthogonal mesh, suggesting that all available septin-septin contacts must be implicated in building the two-layer mesh, even at high septin concentrations.

**Figure 2:**
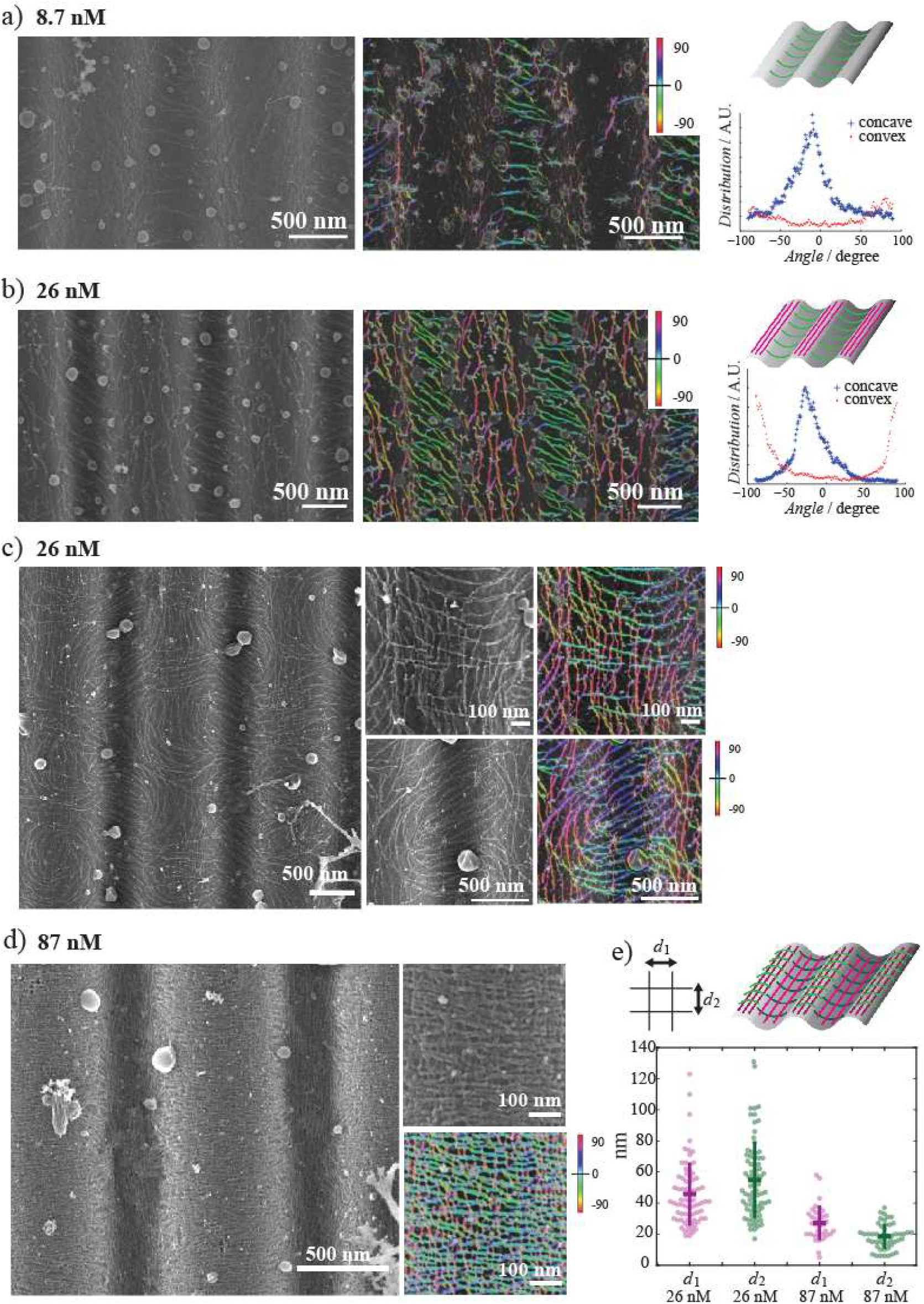
SEM observation of septins filaments on lipid bilayers supported by undulated solid substrates. Undulated solid substrates with 1.6 μm of periodicity and 0.20 μm of amplitude. Samples are prepared with different coucentration of septins dispersed in 10mM Tris buffer containing 75 mM of NaCl. A, B) SEM image of septins filaments prepared at A) 8.7 nM of septins and B) 26 nM of septins in solution, respectively. Raw SEM image is shown on the left. Segmented filaments with color displayed according to the orientation of filaments was overlaid on raw SEM image and is shown in the middle. Distributions of orientation at concave and convex part of substrates are plotted on the right graph. Schematic drawing of representation of obtained results is shown at the right top. C) SEM image of septins filaments at 26nM septins in solution. Image taken covering a wide range is shown on the left, and image focused on mesh structures appeared on convex region and circular pattern of septins filaments on concave region are shown on top right and bottom right, respectively. Segmentation results of filaments with color displaying the orientation of filaments is shown on the right. D) SEM image of septins filaments at 87 nM septins in solution. Image taken covering a wide range is shown on the left, and image focused on mesh structures appeared on convex part of substrates are shown on the right. Segmentation results of filaments with color displaying the orientation of filaments is shown on right bottom. E) Measured distances between two adjacent septins filaments at the center of convex region. Distances *d*_1_ and *d*_2_ correspond to the distances between two adjacent filaments in the direction parallel to the 1D wavy line of the substrate (1^st^ layer of septins on membranes) and perpendicular to the 1D wavy line of the substrate (2^nd^ layer of septins bound to the 1^st^ layer of septins), respectively. Measurement was done depending on the concentration of septins in solution (*d*_1_ = 46±20 (nm) and *d*_2_ = 55±24 (nm) in 26 nM, *d*_1_ = 27±11 (nm) and *d*_2_ = 19±8 (nm) in 87 nM).

**Figure 3.**
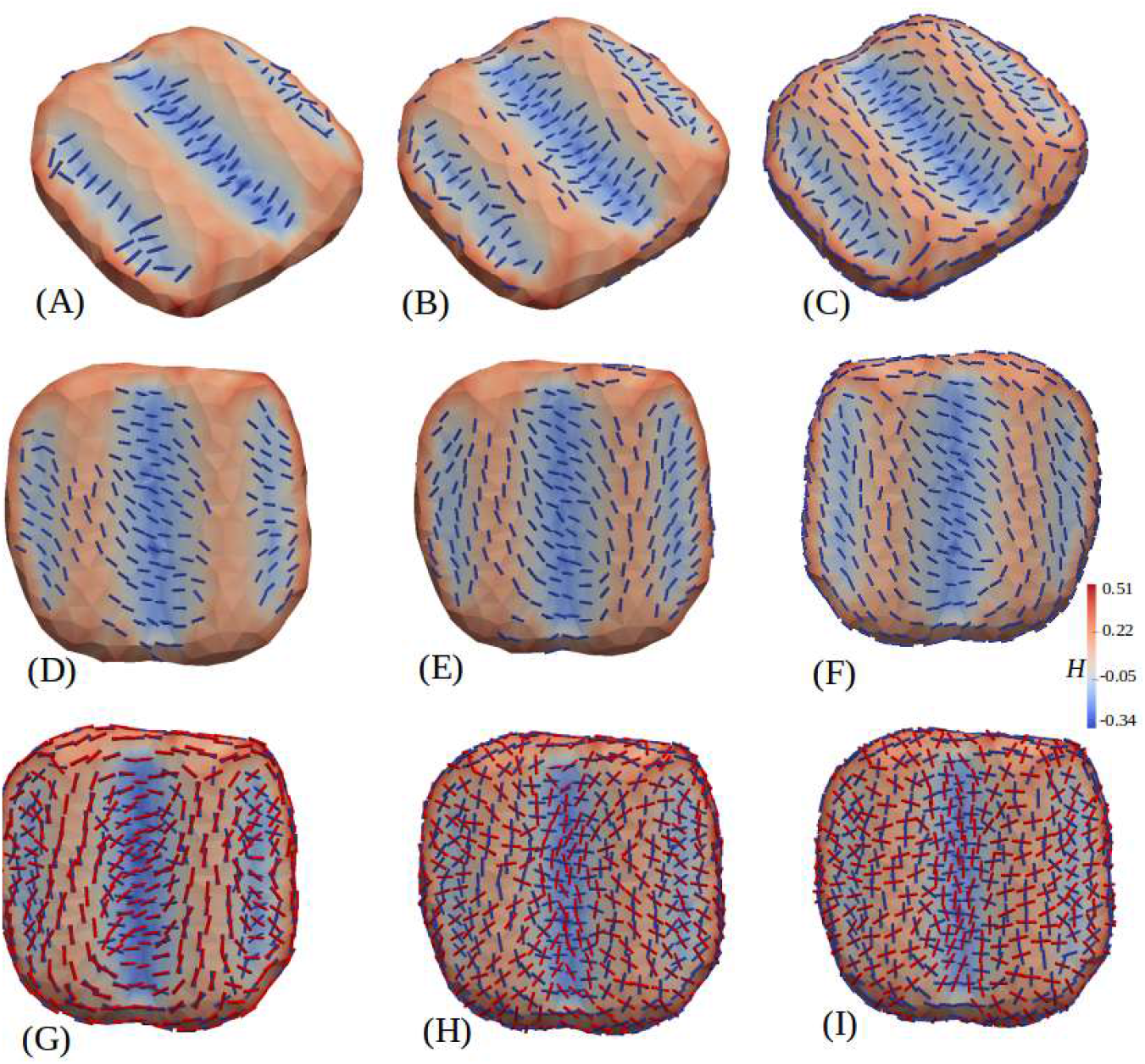
Simulation on wavy patterned surface. Septin on rigid wavy substrate from MC simulation of our model (Eq.1), with septin’s intrinsic curvature *C*_∥_ = –1. The color bar represents local mean curvature of the substrate. The panel (A,B,C) has one layer of septin, while panel (G,H,I) has two layers of septin. Reduction of intrinsic curvature (*C*_∥_ = –0.3 c_∥_) in (D,E,F) changes the septin alignment in the valleys (blue) towards larger radius of curvature, oblique to the direction of undulation. The septin on the hills (red) also turn and form circular structures along with the ones in the valley, see (F). Both in the 1st and 2nd row the population of septins are increased gradually from left toright. In (G,H,I) the strength of interaction (ϵ) between the two layers of septin are increased from left to right (*ϵ* = 1,3,5 and 6). In (I), with *ϵ* = 6, the orientations in the two layers become fully perpendicular. Here the other parameters are *κ* = 20, *κ*_∥_ = 25, *κ*_⊥_ = 25 and *ϵ_LL_* = 2 in K_b_T units and *C*_⊥_ = 0, in arbitrary units. All the parameters are same for both layers of proteins. The color bar indicates local mean curvature of the surface.

### Coarse grained model to capture the organization of septins on wavy substrates

We employed a phenomenological model which had been used earlier to explain how nematic filaments can organize themselves on deformable membranes to reshape them^31,32^. The model is based on the ability of filaments to deform the underlying membrane anisotropically and the tendency of these filaments to arrange parallel to each other forming nematic order. This model had been used for various systems such as GUV coated with FtsZ filaments having intrinsic curvatures, as well as GUVs coated with DNAs displaying no intrinsic curvature^31,33^. Both systems showed the formation of membrane tubes. So far this model had been used to describe the behavior of a single layer of filaments arranged on a deformable membrane^31,32^. In the present work, we have adapted this model to a multilayered system in order to describe how two layers of septin filaments can arrange perpendicular to each other when bound to a non-deformable membrane. We thus modeled the septin coated membrane as two nematic fields adhering to a fluid membrane surface. The first one directly adheres onto the membrane while a second one interacts with both the membrane and the first layer of protein. The details of the model are presented in the method section. The main results, obtained by Monte-Carlo simulations of this model, describing the arrangement of septins on non-deformable wavy surfaces, similar to our wavy curved experimental substrates, are shown in Figure 3, and turn out to be qualitatively consistent with the experimental results. Simulation results in the presence of a single septin layer are presented in Figure 3.A-F. At low density, filaments first populate the troughs, orientating themselves along the curved concave surface with negative curvature, similar to the filament’s own intrinsic curvature, i.e., 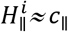 (see Figure 3.A). Here 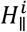 is the local curvature of the surface along the filament length and *c*_∥_ is the intrinsic curvature of the filament. Both 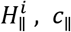, *c*_∥_ being negative here, the energy cost 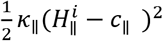 is minimized. Further, at higher density filaments distribute onto the convex part of the wavy surface (the crests) but orient parallel to the channels along the non-curved direction with zero curvature (see Figure 3.B-F) i.e., 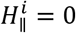, in agreement with Beber et al.^23^. The corresponding energy cost 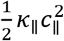 is the minimum, because for any other orientation of the filament on this convex cylindrical shaped surface 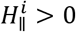, and thus 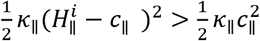. In case the intrinsic curvature of septins was lower than that of the troughs of the wavy substrate, the filaments made an angle with the transverse direction to the channel, (see Figure 3.D-F) seeking out lower 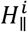 to match with their own intrinsic curvature. This was also pinpointed in our experimental results where filaments were slightly tilted sideways from a pure orthogonal orientation to the wavy undulations (see Figure 2.A and 2.B). At higher septin densities, simulations (see Figure 3.F) presented circular patterns spanning the convex and concave parts of the channels that were similar to the experimental observations (see Figure 2.C). The lower panel (G,H,I) in Figure 3 shows the effect of an additional second layer. In our model, two filaments at the same location but belonging to two different layers, interacted via a repulsive interaction strength *ϵ*, which favored mutually orthogonal orientation between the two filaments (see details in the model section). From left to right, from G to I, *ϵ* was increased. Note that, here, the intrinsic curvatures of filaments did not change in both the first and second layers. For low *ϵ* (see Figure 3.G), both layers tend to align parallel to one another in the channel troughs and crests even at the cost of *ϵ*. However, when *ϵ* dominated over the membrane-septin interaction both layers simultaneously could not minimize their interaction energy with the membrane. At relatively high interaction strength (*ϵ*), the filaments were orthogonal to each other (see Figure 3.I) across the layers.

### Micrometric membrane reshaping by polymerization of human septins octamers

Given that septin filaments specifically arrange on micrometric membrane curvatures (Figure 2–3), one could assume that septin filaments could reshape free deformable membranes. We have thereby examined how septins can deform and reshape the membranes of Giant Unilamellar Vesicles (GUVs) by confocal fluorescence microscopy. To finely describe how septins deform membranes, a phase diagram was built where both the protein and salt concentrations were varied. GUVs containing both DOPS and PI(4,5)P_2_ were prepared and osmolarities within GUVs were carefully adjusted to match the external buffer osmolarity in order to avoid any osmotic shock. Figure 4 presents a typical observation following the incubation of GUVs with human septins octamers-GFP for more than one hour. Without septins, the control GUVs exhibited a rather smooth membrane (Figure 4.B-1). (i) At low bulk septin concentrations (about 1 nM), septins did not distribute homogeneously on membranes (Figure 4.B-2), represented as “weak interaction” in Figure 4.A. Instead septins localized at minor membrane defects (small vesicles or lipid aggregates attached on the surface of GUVs) or at vesicles interconnections (Supplementary. Figure 7.A). When increasing the septin concentration, the surface of septin coverage on GUVs rose (Figure 4.B-3). In ionic strength conditions below 175 mM NaCl, micrometric deformations of GUVs were observed in more than 30% of vesicles, above 10 nM septin concentrations (“intermediate interaction” in Figure 4.A). The deformation rate increased with increasing septin concentrations. At septin concentrations above 100 nM (“strong interaction” in Figure 4.A), more than 50% of the observed vesicles were deformed. Reshaping thus required a high density of septins filaments on membranes. Most of the deformed GUVs displayed micrometric periodic “bumpy” convex shapes, which remained static when imaged for about one minutes. The observed deformations were often irregular (approx. 70%, N_vesicles_ = 62, n experiments = 6), while some “bumps” (approx. 30%) could be regularly distributed and similar in dimensions as shown in the image displayed in Figure 4.B-5 and 4.B-6. The membrane surface was thus covered with convex periodic “bumps” (enlarged in Figure 4.C). We checked that these deformations did not result from the presence of GFP covalently attached to septins. Indeed, “bumpy” GUVs were also observed using dark human septins octamers (Figure 4.B-7). In high salt buffer (above 250mM NaCl, in conditions where the polymerization of septins is inhibited), at any septin concentrations, most of GUVs did not undergo membrane reshaping although GUVs are fully coated with septins. Therefore, membrane reshaping is correlated with the polymerization of septins octamers into filaments. The observed bumpy deformations in deformed vesicles displayed a 2-6 μm periodicity (3.5 ± 2.2 μm (mean ± SD), median = 2.8 μm, N= 18 vesicles) (Figure 4.C), which is in good agreement with the periodicity of membrane deformations induced by yeast septin octamers (varying from 2 to 6 μm)^23^. However, the curvature of the deformations (convex here) was opposite to this imposed by budding yeast septins (concave).

**Figure 4:**
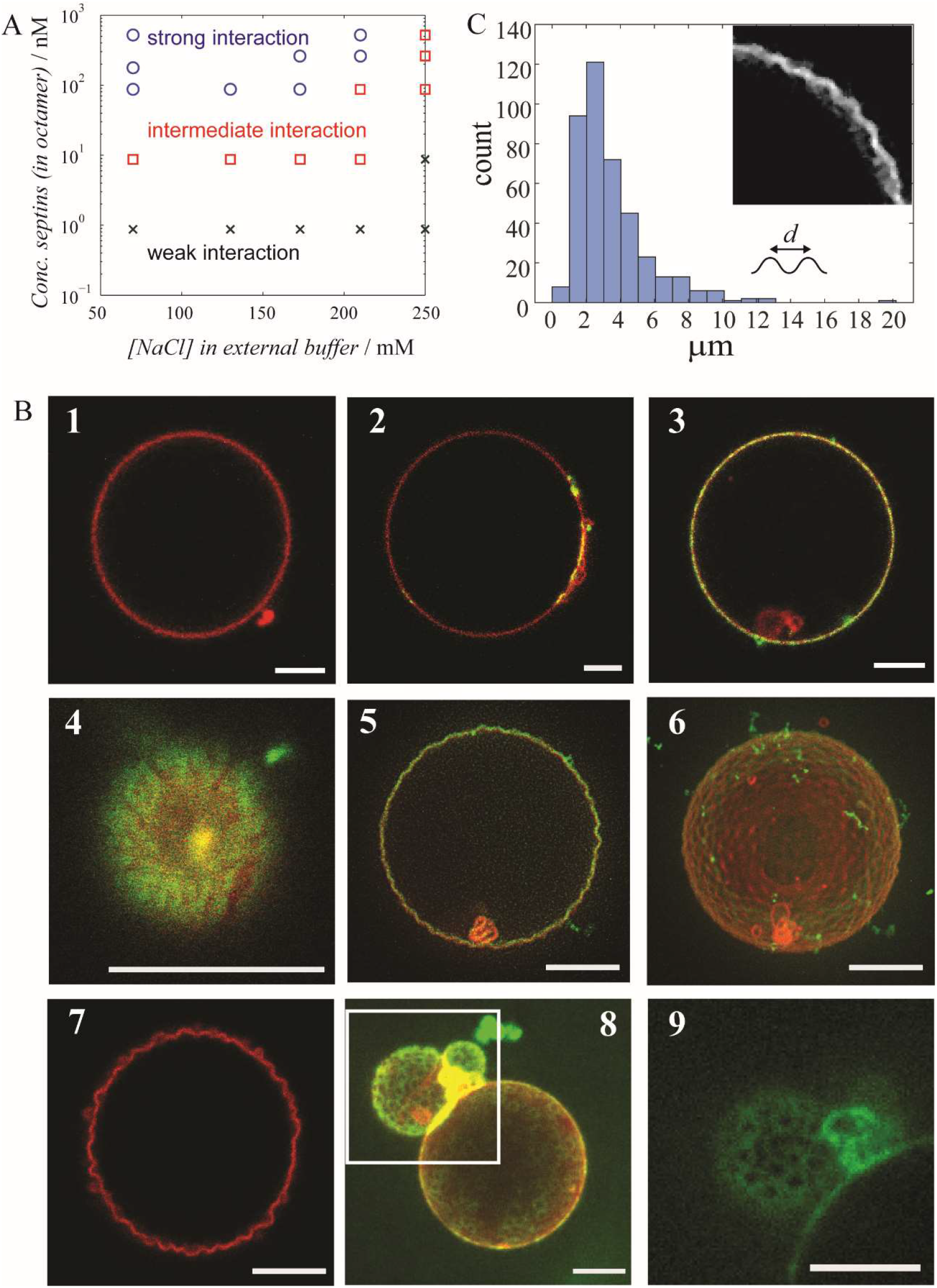
Characterization of the interaction between human septins and membrane. A-Phase diagram of GUV behavior as a function of concentration of NaCl in external buffer and concentration of human septins octamers (in octamer). B-Confocal fluorescence microscopy images of slice of the GUVs. Lipids and septins are both visualized as red and green, respectively, and those signals are overlaid on images. (1) A GUV without human septins octamers in 10mM Tris pH7.8 buffer containing 70 mM NaCl. (2) An example image of GUV found in the region “weak interaction”. GUV is with 0.87nM septins in 10mM Tris buffer contiaining 70mM NaCl. (3, 4) Example of GUVs partially covered by septins. GUVs were incubated with 4.4nM septins in 10 mM Tris buffer containing 70 mM of NaCl. Image 4 is a two-dimensional image and recorded at the bottom of the surface of a GUV. (5, 6) Deformed GUVs found in the region “strong interaction” in phase diagram. Two-dimensional and three-dimensional reconstituted images of GUVs with 176nM septins in 10 mM Tris buffer containing 70 mM NaCl. (7) A deformed GUV with 36 nM of dark septins in 10 mM Tris buffer containing 70mM NaCl. (8, 9) Ring organization of septins bound to membranes at high salt condition. An image of three-dimensionally reconstituted GUV interacting with 520nM septins in 10 mM Tris containing 250 mM NaCl. Image 9 is a two dimensional image of septins (green channel) zoomed at the region of square in image 8. Scale bars = 5 μm. C) Distance between peak of bumps in deformed GUVs (N= 18 vesicles). Different concentrations of septins are plotted on the graph. Each one gives similar distributions (see Supplementary. Figure 8).

In non-polymerizing conditions (> 250 mM salt concentration) and at high septin concentration (> 260 mM), septins organized into rings. Their diameters were measured at 0.96 ± 0.13 μm (N_vesicle_ = 11) as shown in Figure 4.B-8 and Figure 4.B-9. At such ionic strengths, electrostatic interactions should be screened. Without PI(4,5)P_2_ and in the presence of charged DOPS only, those rings did not self-assemble anymore (see Supplementary Figure 7.C and 7.D). The requirement of PI(4,5)P_2_, at high salt, suggested that the septins-membrane interaction in high salt conditions was driven by non-electrostatic specific septin-PI(4,5)P_2_ interactions.

We have demonstrated here that human septin complexes do deform GUVs in a periodic fashion with periodicities similar to the one imposed by our engineered wavy patterns (see Figure 2). The observed mesh-type structure of septins on supported lipid bilayers at high septin concentration probably reflects the septins organization on deformed giant vesicles. Obtained periodicities in GUVs bumpy deformations (2-6 μm) is slightly larger than the periodicity existing in 1D wavy substrates (1.6 μm). Filaments at concave region of supported lipid bilayer are indeed not oriented perfectly perpendicular to the longitudinal axis of 1D periodic wave of substrates, and seem to relax their bindings by tilting from the perfect perpendicular angles.

To better interpret our observations, we used the coarse grained model which was used previously to describe the septin orientation on wavy non deformable substrates (see Figure 3). Except that, in this section, the septins resided on the deformable fluid membrane of a GUV (see method section). Therefore, the filaments-membrane interaction could, in that context, tune the membrane/vesicle shape. We also considered the same repulsive interaction *ϵ* which promotes the local orthogonal orientation between the septins that sit on both layers. Figure 5 shows the results obtained for fluid, deformable surface holding two-layers of nematically organized filaments. The experimental observation showed that membrane deformations frequently occur when septins concentrations are high (Figure 4). Septins were observed to organize themselves in two layers. In the experimental results, the second layer appeared before full saturation of the first layer. This implies that above a given concentration, the septin deposition on the vesicle increases on both the 1st and the 2nd layer because of the increase of septin concentration in the solution. We assumed that the growth of the septin density on the 1st layer weakened the septin-membrane interaction of the 2nd layer, because the membrane surface was dominantly covered with the septins from the 1st layer. Hence, at high concentrations, the effect of the 2^nd^ layer got weaker, while the effect of the 1^st^ layer got stronger. We implemented this by increasing *κ*_∥_ and *κ*_⊥_ of the 1^st^ layer and simultaneously decreasing *κ′*_∥_ and *κ′*_⊥_ of the 2^nd^ layer, with increasing septin concentrations. Based on these hypotheses, the deformation of GUVs could be reproduced from our simulation, as shown in Figure 5. At low concentration, both layers had equal strengths of interaction with the membrane and our simulation showed that for *κ*_∥_ = *κ′*_∥_, all undulations vanished, resulting in a smooth spherical surface. When *κ*_∥_ > *κ′*_∥_ and *κ*_⊥_ > *κ′*_⊥_, as shown in Figure 5.C-E, membrane deformations similar to our experimental observations were obtained. Membrane deformations can thus be dominated by the interaction between the first layer of septin and the membrane. At low concentrations of septin, when *κ*_∥_ and *κ′*_∥_ were estimated to be equal, the membrane deforming ability of the two layers canceled each other, provided *κ*_∥_ and *κ′*_∥_ were small in magnitude relative to *ϵ*, ensuring locally orthogonal alignment of septins in the two layers. This was a nontrivial effect because, at a given point on the membrane, one could consider an isotropic concave valley (i.e., part of an inverted hemi-spherical bowl) which could accommodate two septin filaments that were mutually orthogonal and belonging to the different layers. However, in a volume preserving scenario (which corresponded to our conditions) valleys would also be accompanied by hills where the surface curvature was positive and therefore unfavorable to filaments (since *c*_∥_ < 0), thus causing a positive energy. Here, we note that it is indeed the case when intrinsic curvatures of filaments are identical in the first and second layers. In a real situation, septins can be oriented differently at first and second layer, and it can change the effect of the second layer on membrane deformations. In addition, surface tension also opposes surface undulations. It turns out that instead of growing such valleys and hills the vesicle prefers a smooth surface which has a low positive energy cost. An approximate energy comparison is given in supplementary text 1.

**Figure 5.**
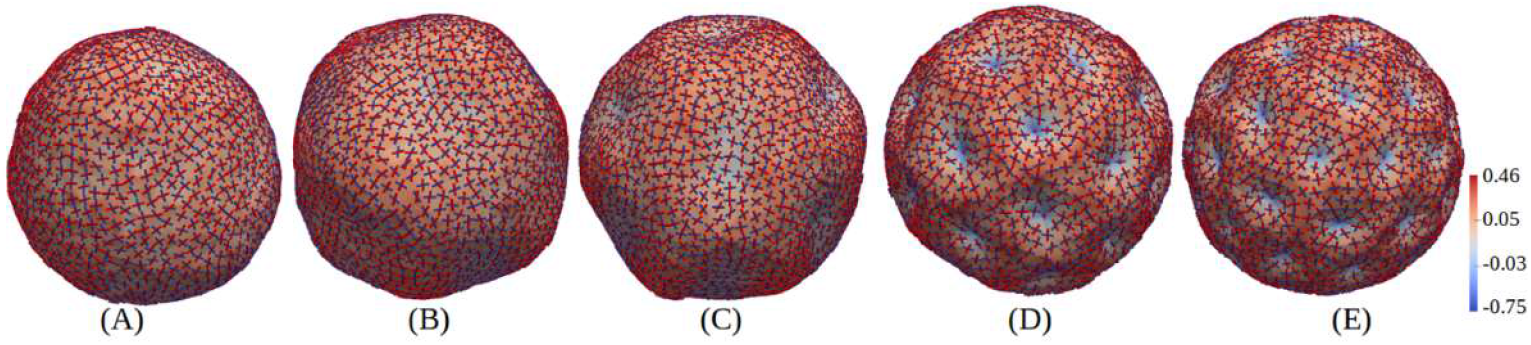
Simulation describing membrane deformations by nematically ordered filaments. Septin on deformable vesicle. The ratio of the corresponding induced bending rigidities for both layers 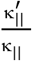 and 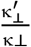 are 1 (in A), 0.5 (in B), 0.1 (in C), and 0 (in D,E). Corresponding *κ*_∥_ and *κ*_⊥_ here are 5, 10, 15, 25 and 40, for A-E respectively. Here, the other parameters are *κ* = 20, *ϵ_LL_* = 1 and *ϵ* = 1 (in K_b_T units), *C*_∥_ = –1 and *C*_⊥_ = 0. All the parameters are same for both layers of proteins. The color bar indicates local mean curvature of the surface.

## Discussion

After investigating the behavior of budding yeast septins bound to membrane, clues from observations in solution^4,26^ and septin localization in vivo^34^ indicated that the metazoan and thereby human septin interaction with lipids would bring unexpected and novel outcome, as compared with fungi septins. In solution, human septin octameric complexes self-assemble similarly to Drosophila septins^35^. Those metazoan octameric septin complexes spontaneously bundle through lateral filamentous interactions^4,26,35–38^ and appear as straight spine-like or circular bundles of paired or single filaments (Supplementary. Figure 9). On membranes Drosophila septins form tight sheets of parallel filaments^4^, similarly to budding yeast septins. On another hand, thoroughly studied budding yeast septins assemble into stable paired filaments and only occasionally bundle^36,39^. S. Cerevisiae and homo sapiens septins thus display different behaviors, when it comes to self-assembly. Even though septins have evolved from an ancient ancestor^40^, phylogenetic analysis have shown that within the seven subgroups in which septins are classified, only one group (1A) encloses both budding yeast Cdc10 and human septin Sept9. Besides, human septins and particularly Sept9 can be expressed in a variety of splice variants which contrasts with fungi septins^14^.

We have therefore used a combination of methods to follow the interaction and curvature preference of human septins bound to liposomes at different scales (ranging from a few tens of nm to μm). We analyzed the organization of the human septin octameric complex interacting with wavy substrates of varying micrometer curvatures, and coated with supported lipid bilayers. Using electron and fluorescence microscopies, we were able to describe the features of our samples at different scales (from nanometer to micrometer ranges).

In the presence of a biomimetic membrane (supported lipid bilayer, lipid monolayer), we propose in the present work that the septin-membrane interaction prevails over the septin-septin lateral association, preventing the self-association of thick bundles. Single or paired filaments thus assemble onto membranes without bundling. We found that human septins systematically assemble into arrays of orthogonal filaments contrary to budding yeast septins that preferentially self-assemble on membranes as parallel filamentous arrays and would only occasionally arrange as networks of filaments^2,23^. Moreover, orthogonal gauzes of human septin filaments are consistently observed on liposomes (Supplementary Figure 2) or on supported lipid bilayers (Figure 2.D). The inter-filament distance (around 30 nm), extracted from the SEM on undulated substrate, agree either with the spacing of paired filaments or with an octameric periodicity. At the highest densities, the first layer of septin filaments directly apposed onto the lipids display a regular spacing of about 30 nm, in agreement with a septin octameric length. The second septin layer sits orthogonal to the first layer. Within the second layer, our analysis tends to show that filaments are closer from one another (from 10 to 20 nm) and would thus coincide with filament pairing mediated by coiled coils. To induce this network, the first septin should be specifically arranged in a parallel fashion and create a scaffold to recruit the second perpendicular layer.

Based on the molecular specificities of septins, Figure 6 displays a molecular model that would account for our observations. X-rays crystallography studies^41^ suggest the versatility of the possible septin-septin interactions mediated through the coiled coil domains of Sept6, Sept7 and Sept2. Antiparallel, parallel arrangements of Sept2-Sept2 coiled coils as well as homo dimeric or hetero dimeric, parallel or antiparallel coiled coils involving Sept6 and Sept7 can be generated. In our samples, the lateral organization between adjacent filaments in the first layer is probably triggered by the pairing of filaments using CCs mediation (most likely Antiparallel coiled coils involving SEPT6-7^42^. The filament side interacting directly with the membrane could mediate its interaction through the alpha 0 domain of septins (PB1). Leonardo et al.^42^ suggest that the space between the CC and the membrane could integrate a chain of hydrogen bonds because of the limited rotation of the CC superhelix thus displaying a hydrophilic side towards the membrane. The septin-membrane interaction would thus be stabilized through the coiled coils. The presence of polybasic regions at the very end of SEPT6 and SEPT7 C-termini also strengthen the septin-membrane interaction. The first septin layer might, in turn, interact with the second layer, possibly from Sept2-Sept2 antiparallel coiled coil interactions, one helix from each layer (see Figure 6). The second layer would thus stabilize the first septin layer and induce an octameric repeat in between the septin filaments from the first layer. The higher density of the second septin layer might impose a higher ordering of the first layer. Short distances (from 10 nm to 20 nm) are indeed observed between filaments in second layer. It would be possible that two filaments are closely paired and juxtaposed when interacting with the first septin layer or connected via coiled coils. Besides, we believe that coiled coils pairing remains rather flexible and induce both the measured dispersity in the inter-filament distances and enough flexibility to allow, to some extent, the septin filaments adaptation to curvatures. Finally, only two layers of filaments can be observed. Septins thus do not accumulate into successive layers, suggesting that the coiled coils that possibly mediate the interaction are all already engaged in the septin membrane and septin-septin interaction. A similar behavior had already been observed in the Drosophila septins organization onto membranes^4^.

**Figure 6.**
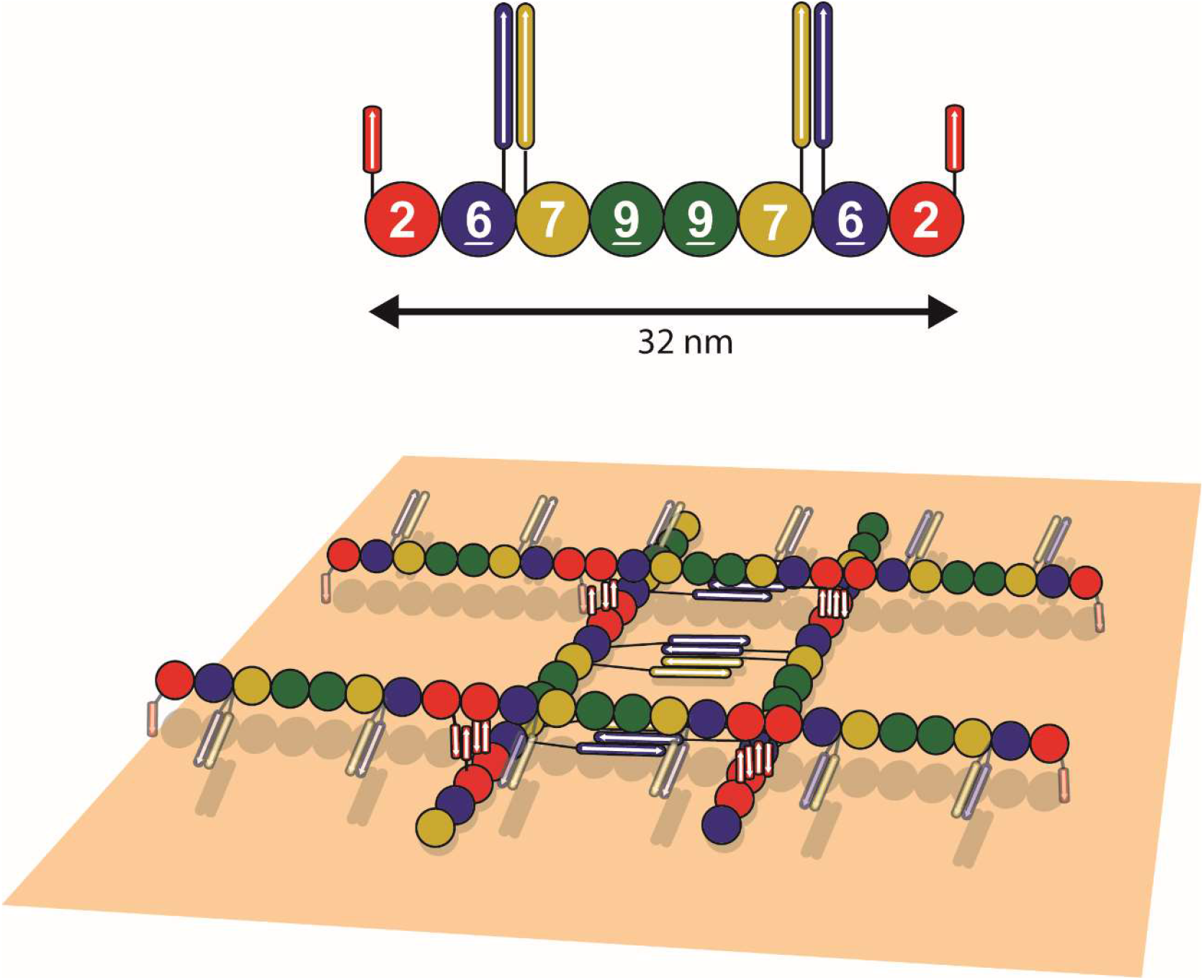
Model of orthogonal network assembly. Schematic drawing of octameric septins (top) and a possible model of orthogonal network array of human septins filaments. Note that the orientation of parallel coiled coils in the second (upper) layer of septins could diverge from this proposal in terms of orientation.

The curvature sensitivity of septins at the micrometer scale has been first highlighted in fungi in vitro and has been discovered and intensively investigated in the budding yeast model^17,23,28^. Given that human septins organize in a more complex network-like manner, this model needs to be revisited. In mammalian cells, septins also interact with membranes at specific locations where micrometric curvatures are present, both positive and negative^11,19,43^. The specificity of septins to micrometric membrane curvatures is probably firmly inherited over species. Since septins are involved in membrane remodeling processes, and particularly in human, revealing the mechanisms of membrane curvature sensitivity at the molecular level is key to understand the mechanisms of cellular functions involving human septins.

From the discrepancies observed between human and yeast septin ultrastructural organization, it is thus not surprising to report distinct behaviors for membrane reshaping and curvature sensitivity. In budding yeast, a dense parallel set of septin filaments would deform GUVs in vitro to generate negative micrometric curvatures. Those concave deformations and correlated curvature sensitivity imposed by budding yeast septins were modelled using an enhanced membrane affinity energy with the filaments bent negatively^23^. For human septins, the behavior, in terms of curvature sensitivity, is similar to that of budding yeast septins at low concentration. At low septin concentrations, where the filament density is not sufficient to induce membrane deformations, human septins do not assemble into orthogonal arrays. Instead, they organize as individual filaments (See Supplementary Figure 5.A) and bend negatively on concave geometries while they remain unbent when encountering a hilly positively curved domain. Conversely, at higher protein concentrations, orthogonal gauzes are visualized on both positively and negatively curved domains with similar densities. Our Monte Carlo simulations which consider septins as nematically ordered filaments, closely reproduce those experimental observations. The transition from a low filament density to a dense network of filaments occurs within a short range of concentrations (30-40 nM). We were actually unable to detect any intermediate state in terms of protein density on supported lipid bilayers, suggesting that a cooperative effect tends to facilitate the formation of networks. Those network-like organization most likely reflects the organization of human septins when bound to GUVs. Indeed, on GUVs, the observed deformations follow a curvature which dimensions correspond to those of our wavy substrates. A dense orthogonal network of human septins thus generate positively curved deformations, as also reflected by the simulations (Figure 5). We can thus assume that the formation of orthogonal network of filaments is key to bolster the emergence of membrane deformations in the scope of our current report.

Tanaka-Takiguchi et al.^24^ had captured outwards tubulations events (thereby concave deformations) on GUVs incubated with either brain extract or hexameric human septin complexes (Sept2-Sept6-Sept7). Such micrometric GUV deformations induced by the collective behavior of nematic-ordered filaments has been also discussed in other reports ^31,32^. For example, similar GUV deformations, where GUVs can be deformed and wrinkled before tubulation occurs, were observed using DNA Origami constructs capable of self-assembling into filamentous structures^33^. Franquelim et al. thereby propose that the polymerization of filaments bound to membranes can control membrane reshaping. Our Monte-Carlo simulation has shown (Figure 5) that negative intrinsic curvature of septin (i.e., concave) is able to produce both convex hills and concave valleys on GUV surface. The first layer of septin (blue in Figure 5), which is dominant, orient on the valleys in a vortex like pattern exploiting the concave shape of the valley. On the convex portions of the vesicle, between two valleys, septin filaments orient along the direction which has minimal positive curvature. This is identical to their orientation on the wavy substrate. The second layer of septin (red in Figure 5), which has a relatively weak effect on vesicle deformation, orient along the downhill concave direction towards the bottom of the valley, satisfying its own negative intrinsic curvature. We found that the valleys have the shape of inverted hemispheres rather than pointed inverted cones with reducing apex angle. This energetically benefits both the layers of septin as they attain preferred negative curvature. On the convex hills both layers pay energetic price.

Kumar et al.^31,44^ have pointed out from their Monte Carlo simulations that the intrinsic curvatures of filaments imposing anisotropic membrane curvatures (both longitudinal and orthogonal to the direction of the filaments) can drive either concave or convex deformations of liposomes, with the filaments ordered in a nematic fashion. We have specifically chosen negative intrinsic curvature for human septin filaments along their length because in Yeast, also, they were found to have negative intrinsic curvature of similar magnitude. However, yeast septin showed additional bundling interaction which is absent for human septins. This general model of nematic filaments interacting with GUVs, imposing anisotropic membrane curvature, has been relevant for other systems as well. For example, artificially tuning the intrinsic curvature of FtsZ can trigger the emergence of either concave or convex deformations and tubulations on vesicles^45^. However membrane curvature and vesicle tubulation can also be induced by chiral nematics having no intrinsic curvature or bundling interaction^46^. Overall, application of this model to human septins help us rationalize our experimental observations of the orientation pattern of septins on rigid wavy surfaces as well the nature of shape deformations of vesicles.

## Conclusion

We studied, *in vitro*, the organization of human septins octamers (SEPT2-GFP, SEPT6, SEPT7, SEPT9i1) on reconstituted lipid model membranes. By electron microscopy, it was revealed that human septins filaments assemble into a perpendicular mesh-like network on membranes constituted with two layers of nematically ordered septins filaments perpendicularly bound to each other. Human septins are able to reshape soft deformable membranes (Giant unilamellar vesicles). On undulated supported lipid bilayer with micrometric periodicities similar to the scale of those deformations, the filaments orient according to the sign of local curvatures, resulting in regular mesh structure in distances and angles. As compared with septins from other species (yeast and fly), human septins organize in a more complex network-like manner. Our Monte Carlo simulations could recapitulate the behavior of human septins considering the filaments as rod like nematic liquid crystalline objects.

## Materials and Methods

### Chemicals

Common reagents (ethanol, acetone, chloroform, sucrose, sodium chloride, Tris) were purchased from VWR reagents and Sigma-Aldrich Co.. L-α-phosphatidylcholine (EPC, 840051P), cholesterol (700000P), 1,2-dioleoyl-sn-glycero-3-phosphoethanolamine (DOPE, 850725P), 1,2-dioleoyl-sn-glycero-3-phospho-L-serine (DOPS, 840035P), and L-α-phosphatidylinositol-4,5-bisphosphate (PI(4,5)P_2_, 840046P) were purchased from avanti polar. Bodipy-TR-ceramide was purchased from Invitrogen (D-7540).

### Protein purification

Human septins octameric complexes containing SEPT2-GFP, SEPT6, SEPT7, SEPT9i1 were co-expressed in *Escherichia coli* and purified as described in detail elsewhere^47^. Briefly, septins are purified by nickel-histidine, streptavidin-biotin affinities and ion exchange chromatography. septins complex mostly consists of mixtures of octamers SEPT2GFP-SEPT6-SEPT7-SEPT9-SEPT9-SEPT7-SEPT6-SEPT2GFP. Those complexes are conserved in highly salted solutions (ca. 300mM KCl, 50mM Tris pH 8) at −80 °C (after quick freeze in liquid nitrogen) to avoid polymerization and aggregation of complexes in solution.

### GUV assay

Giant unilamellar vesicles (GUV) were prepared by either platinum wire method^27^ or PVA-gel assisted methods^27^ with lipid mixtures of EggPC 56.5%, Cholesterol 15%, DOPE 10%, DOPS 10%, PI(4,5)P_2_ 8% and Bodipy TR ceramide 0.5%. In both methods, lipids were resuspended in 10 mM Tris pH 7.8, 50mM NaCl and desired concentration of sucrose. Formed GUVs were collected and transferred in solution with 10mM Tris pH 7.8 containing desired concentration of NaCl. Osmolarity inside and outside of vesicles were adjusted by the amount of sucrose according to the NaCl concentration of external solution. Septins were added to GUVs from outside, and samples were incubated for more than 1 hour. Samples were then transferred to the glass chambers passivated with 5wt% ß-Casein/10mM Tris pH 7.8 75mM NaCl and were observed under confocal fluorescence microscopy. Confocal experiments were performed on a Nikon Eclipse TE2000 inverted microscope equipped with software EZ-C1, and images were analyzed by imageJ software. To obtain three-dimensional images, Inverted Eclipse Ti-E (Nikon) confocal microscope equipped with Spinning disk CSU-X1 (Yokogawa) and Live-SR (Gataca Systems) integrated in Metamorph software was used. Images are taken by Prime 95B (Photometrics) and are further analyzed using imageJ software.

### Cryo-electron microscopy and cryo-electron tomography

Lipid mixture (EggPC 57%, Cholesterol 15%, DOPE 10%, DOPS 10%, PI(4,5)P_2_ 8%) dissolved in chloroform was quickly dried under nitrogen. Obtained lipid dried films are further dried under vacuum for more than 30 min. Lipids are resuspended in 10mM Tris pH 7.8 buffer containing 75mM NaCl by voltex to obtain various size of LUVs. LUVs suspension was diluted and septins were added to a final concentration of 17 nM septin and 0.0125-0.025 g/L lipid. Mixture was incubated for 1h, and 10 nm size gold beads were added to the solution just before the deposition of electron microscopy grid. 4 μL of solution was deposited on a glow discharged lacey carbon electron microscopy grid (Ted Pella, USA). Most of the solution was blotted away from the grid to leave a thin (<100 nm) film of aqueous solution. The blotting was carried out on the opposite side from the liquid drop and plunge-frozen in liquid ethane at –181 °C using an automated freeze plunging apparatus (EMGP, Leica, Germany). The samples were kept in liquid nitrogen and imaged using a Tecnai G2 (FEI, Eindhoven, Netherlands) microscope operated at 200 kV and equipped with a 4kx4k CMOS camera (F416, TVIPS). For cryo-electron tomography, tilt series were collected in low dose mode, every two degrees, 0 to −34°, then +2 to 60° and finally −36 to −60° to minimize irradiation at the lowest angles. The dose per image was set around 1 electron per Å^2^. The imaging was performed at a magnification of 50,000. The consecutive images were aligned using the IMOD software with a help of gold beads. SIRT reconstruction was carried out using for volumetric reconstitution. The segmentation was performed manually using IMOD.

### Scanning electron microscopy

Wrinkled poly(dimethyl siloxane) PDMS substrates having 1.6±0.1 μm of period and 0.20±0.05 μm of amplitude were designed and fabricated to achieve pattern curvature of ±3 μm^-1^ as described by previous report for fabrication of 1D wave^48^. Fabricated wavy PDMS substrates was used to make a surface replica using UV-curable adhesive (Norland Optical Adhesive NOA71). Small drop (ca. 5 μL) of NOA71 was placed on circular cover slip (12 mm diameter) and wavy surface of PDMS chip had a contact with NOA71. NOA71 was cured under UV lamp for 7 min and PDMS chip was carefully pealed and stored for further replication. SUVs solution to prepare supported lipid bilayer on top of wavy substrate was prepared as following. Desired amount of lipid mixture (EggPC 57%, Cholesterol 15%, DOPE 10%, DOPS 10%, PI(4,5)P_2_ 8%) dissolved in chloroform was quickly dried under nitrogen and obtained lipid dried films are further dried under vacuum for more than 30 min. Lipids are resuspended at 5 g/L in 20mM citrate buffer pH4.8 containing 150mM NaCl, and suspension was sonicated for around 30 minutes until the solution became transparent. Obtained SUV suspension was kept in −20 °C freezer, and before the use, it was diluted to 1g/L with the same citrate buffer.

100 μL of 1g/L SUV suspension was deposited on the prepared NOA wavy substrates treated with plasma for 1-3 min in advance. Sample was incubated for 1h, and substrate was washed thoroughly 4 times using 20mM citrate buffer pH4.8 containing 150mM NaCl and then 4 times using 10mM Tris pH 7.8 buffer containing 75mM NaCl. After the wash, septins were added on the substrate with various concentration in 10mM Tris pH 7.8 buffer containing 75mM NaCl. After 1h of incubation, samples were fixed with Glutaraldehyde 2% in sodium cacodylate 0.1M for 15 min. The samples were washed three times with sodium cacodylate 0.1M and incubated for 10 min with a second fixative (Osmium tetroxide 1% in sodium cacodylate 0.1 M). After three washes with water, the samples were incubated with tannic acid 1% for 10 min and subsequently washed three time with water before being incubated with Uranyl acetate (1% in water) for 10 min. The samples were then dehydrated using baths with increasing ethanol concentrations (50, 70, 95, 100%) and processed using a critical point dryer (CPD 300, LEICA). After being mounted on a sample holder, the samples were coated with around 1 nm of either tungsten (ACE 200, LEICA). SEM imaging was performed using a GeminiSEM 500 microscope from Zeiss, Germany. SEM images were segmented using Weka trainable segmentation on ImageJ software and were further analyzed using OrientationJ to determine the orientation of the filaments. To extract the periodicities existing in mesh organization of septins filaments, probability map obtained as a result of segmentation was fourier transformed using FFT on imageJ software, and square root of raw power spectrum obtained by FFT was integrated by angles and was displayed as 1D power spectrum. Spectrum was fitted by Lorentzian function to obtain the distance and FHWH.

### Fluorescence microscopy image analysis: measurement of periodicity in GUV deformation observed by fluorescence microscopy

The fluorescence signals from the membranes of a deformed (regular deformation or partially distorted deformation) GUV were used for the measurement of periodicity in membrane deformation by septins. The image was binarized by thresholding and particle analysis in ImageJ software. The GUV was fitted with an ellipsoid to determine the center of vesicles. The x y coordinates of the vesicle perimeter were then obtained by using Find Edge function in the ImageJ software. Distances from the center to a point (pixel) on the vesicle perimeter was obtained for each point of vesicle perimeter. and the distances were plotted as a function of the angles *θ* (see Supplementary Figure 10). The angles *θ* were converted to the actual distances *d* by *d* = *2πR*((*θ* + 180)/360) × *pixel size* (*μm/pixel*). Here, the mean radius *R* was obtained according to the major and minor length of fitted ellipsoid by (major + minor)/2. Local maxima in *l* were found under the condition where *d* > 0.25 μm and l > 0.6 μm). Several images were subjected to this image analysis, and all the distances d were plotted in a histogram.

### SEM Image Analysis: display the orientation profiles on septins filaments

Raw SEM images are segmented by using machine-learning based Weka trainable segmentation in ImageJ software. The whole image was then classified into three categories (septin filaments, small vesicles bound on membranes, and background (supported lipid bilayer only)) (an example shown in Supplementary Figure 11). A probability map was generated by segmentation (see Supplementary Figure 11). The resulting probability map for septin filaments was used for further image analysis. The probability map of septins filaments was first rotated so that the 1D periodic waves of substrates orient perpendicularly. Image J plugin “Orientation J” displayed the orientation of the filaments on probability maps by color, and a spectrum displayed the angle distribution.

### SEM Image Analysis: measurement of the periodicity in septins mesh organization using 2D FFT signals

Raw SEM images were segmented and the probability maps of septin filaments were obtained as described above. From the whole probability map, sections where meshes are clearly visualized on convex region (positive curvature) were cropped and then Fourier transformed by 2D Fast Fourier Transform (FFT) function using the ImageJ software. The resulting 2D FFT signal was radially integrated by using the ImageJ plugin “Radial_Profile_Angle”. The1D spectra of the signals were then plotted in reciprocal nanometer (nm^-1^) as shown in Sup. Figure 4. Several images were subjected to this analysis, and all the obtained 1D spectra were averaged by different positions assuming that the density of septin meshes are uniform over the sample.

### Model for Monte Carlo simulations

The local orientation of the nematic field, which models filament orientation, is denoted by the unit vector 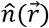 which lies in the local tangent plane of the membrane and is free to rotate in this plane. The vesicle is modeled by a closed, triangulated network of vertices (i = 1, 2,…, N), see Supplementary Figure 6, and filament-membrane interactions are modelled as anisotropic spontaneous curvatures of the membrane, in the vicinity of the filament. Two nematic fields, 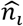 and 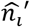, correspond to the nematic field of 1^st^ layer and 2^nd^ layer, respectively, are at the same vertex ‘i’. Two fields prefer to orient perpendicular to each other via an interaction term 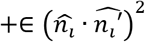 which preserves 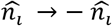 symmetry of the nematics. Nematics at the same layer tend to align with each other via the nearest neighbor nematic interaction energy, commonly modeled by the Lebwohl-Lasher form^49^ 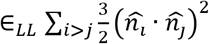. Here *∈_LL_* is the strength of the nematic interaction, in one constant approximation (i.e., when splay and bend moduli are assumed to have the same value). The total energy functional is given by,

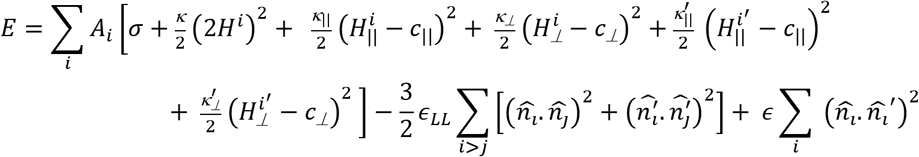

All the terms in the first set of square brackets are summed over surface elements *A_i_* associated with the i-th vertex. The 1st term represents surface tension, and the 2nd is the Helfrich elastic energy^50^ for membranes with isotropic bending rigidity *κ* and membrane mean curvature *H* = (*c*_1_, + *c*_2_)/2. Here, *c*_1_ and *c*_2_ are the local principal curvatures on the membrane surface along tangent vectors 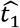 and 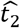, respectively. *κ*_∥_ and *κ*_⊥_ are the induced membrane bending rigidities, *c*_∥_ and *c*_±_ are the induced intrinsic curvatures along and perpendicular to 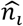 (or 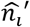 in the second layer) which are the orientations of the filaments in the local tangent planes in the two layers, respectively. Here, for human septins exhibiting intrinsic negative curvature, we set *c*_∥_ to a non zero negative value and *c*_⊥_ = 0. The local membrane curvature along 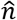 (and 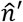, in the second layer)) is given by *H*_∥_ = *c*_1_, cos^2^ *φ* – *c*_2_ sin^2^ *φ* (and *H′*_∥_ = *c*_1_, cos^2^ *φ′* – *c*_2_ sin^2^ *φ′*) and perpendicular to 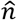 (and 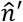) is given by *H*_⊥_ = *c*_1_ cos^2^ *φ* – *c*_2_ sin^2^ *φ* (and 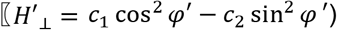, where *φ* and *φ′* denote the angles between the principal direction 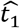 and the filament orientations 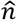 and 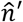, respectively. Note that at a given point on the membrane the principal curvatures *c*_1_, *c*_2_ and the corresponding eigen vectors 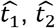, are unique, but the curvatures and *H*_∥_ and *H*’_∥_, along the filaments 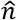 and 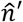, respectively, are different.

In a model for a fixed surface, the local mean curvature *H^i^* at any point on it is predetermined, but the filaments seek out appropriate 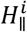 (and 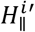 for the second layer) at any point, by changing its orientation, such that it minimizes the global energy of the system. In a model for a deformable GUV, on the other hand, the vertices are continuously moved during Monte-Carlo steps, in order to explore the shape space of the deformable fluid surface. In our simulation the septin concentration is limited by the number of vertices in the network (maximum one in each layer). Also, our nematic interaction acts only between neighboring vertices and energy is lowered if the neighbors are parallel to each other. Therefore there is a tendency in the nematics to self-assemble even if the vertices are sparsely populated at the starting of the simulation. However, in reality, increased septin concentration in the solution decrease the inter-septin distance on the membrane. We model the increased concentration by increasing the strength of the septin-membrane interaction (*κ*_∥_).

## Acknowledgement

We thank Patricia Bassereau and Daniel Lévy for useful advice and discussions. We thank Manos Mavrakis for the kind gifts of the plasmids expressing human septins. This work benefited from the support of the ANR (Agence Nationale de la Recherche) for funding the project “SEPTIME”, ANR-13-JSV8-0002-01 and the project “SEPTSCORT”, ANR-20-CE11-0014-01. B. Chauvin is funded by the Ecole Doctorale “ED564: Physique en Ile de France” and Fondation pour la Recherche Médicale. K. Nakazawa was supported by Sorbonne Université (AAP Emergence). We thank the Labex Cell(n)Scale (ANR-11-LABX0038) and to Paris Sciences et Lettres (ANR-10-IDEX-0001-02). We thank the Cell and Tissue Imaging (PICT-IBiSA), Institut Curie, member of the French National Research Infrastructure France-BioImaging (ANR10-INBS-04).

## Conflict of interest

The authors declare no conflict of interest.

## Notes

### Competing Interest Statement

The authors have declared no competing interest.

### Summary of Updates

Supplementary material has been added

