## supplemental material for "A specific mesh-like organization of human septin octameric complex drives membrane reshaping and curvature sensitivity"

*correspondings authors

[#]Equal contribution of those authors

**Supplementary Movie 1**

Slices within the cryo-tomogram from Figure 1.C displaying septin filaments bound to a liposome. The layers of filaments are visualized on different slices at t=2-4 seconds and t=29-31 seconds. Scale bar:100nm.

**Supplementary Figure 1**

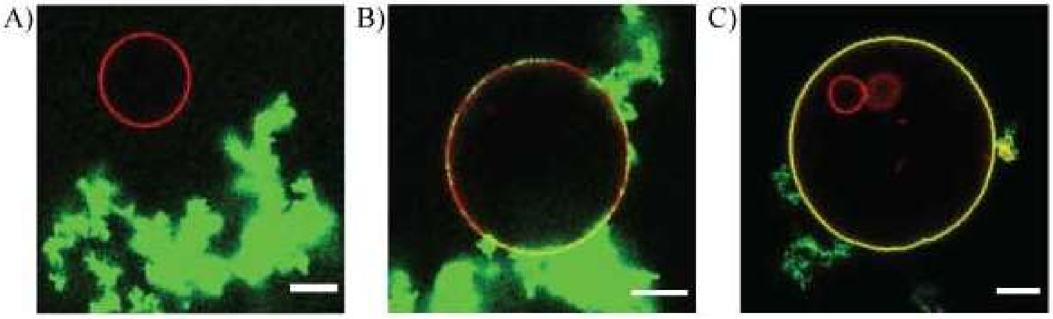


Optimization of the septin/membrane interactions

(A) Typical confocal microscopy image showing the Interaction of GUV without any negatively charged lipids (lipid composition of GUV: EggPC 99.5%, Bodipy TR ceramide 0.5%) in 10 mM Tris pH 7.8 buffer containing 70mM of NaCl. Concentration of human septins octamers-GFP in solution outside GUVs is 260 nM.

(B) Typical confocal microscopy image of GUV containing DOPS and without PI(4,5)$P_2$ lipids (lipid composition of GUV: EggPC 64,5%, Cholesterol 15%, DOPS 10%, DOPE 10%, Bodipy TR ceramide 0.5%) in the presence of 70 mM NaCl, Tris 10 mM pH 7.8. Concentration of human septins octamers-GFP in solution outside GUV is 260 nM.

(C) Typical confocal microscopy image of GUV containing both DOPS and PI(4,5)$P_2$ lipids (lipid composition of GUV: EggPC 56,5%, Cholesterol 15%, DOPS 10%, DOPE 10%, PI(4,5)$P_2$ 8%, Bodipy TR cermide 0.5%) in the presence of 70 mM NaCl, Tris 10 mM pH 7.8. Concentration of human septins octamers-GFP in solution outside GUV is 260 nM.

Scale bars: 5 μm

**Supplementary figure 2**

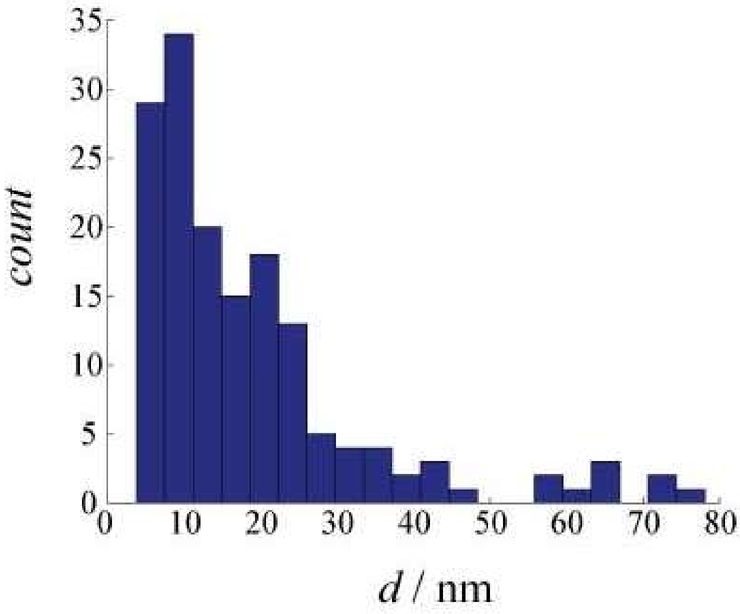


Histogram of measured distances of adjacent septins filaments $d$ = 18 ±15 nm (Mean + SD) observed on 4 different tomograms of LUVs (157 counts, $N_{vesicle}$ = 4). Measurement was done with the segmented images.

**Supplementary Figure 3**

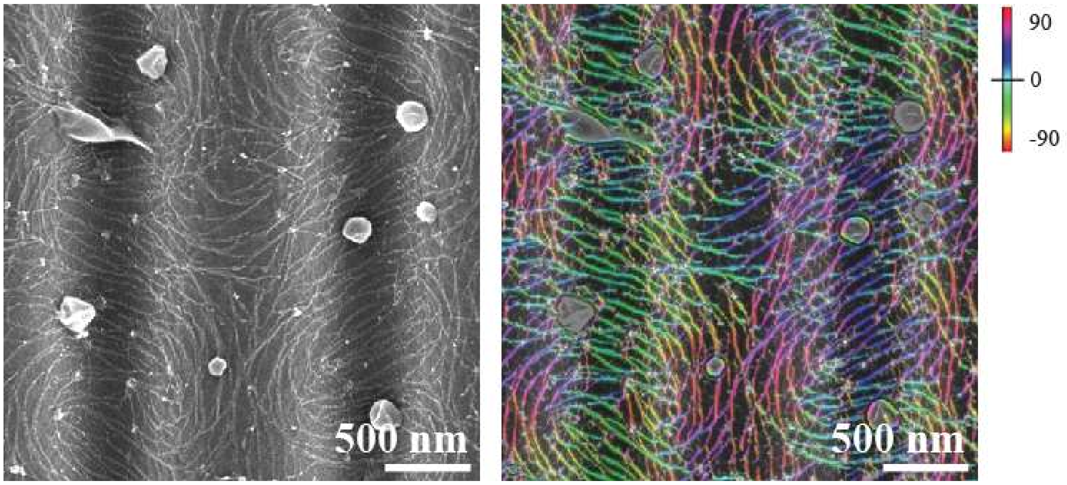


Images for SEM observation of septins filaments on lipid bilayer supported by undulated solid substrates with 1.6 μm of periodicity and 0.20 μm of amplitude. Septins concentration is 26 nM in octamer. Raw image is shown on the left. On the right image, segmented septins filaments by using Trainable Weka Segmentation in ImageJ with color map showing the orientation of those filaments obtained by OrientationJ in ImageJ is superimposed to the Raw image. Spiral organization of septins filaments centered at concave regions are observed.

**Supplementary Figure 4**

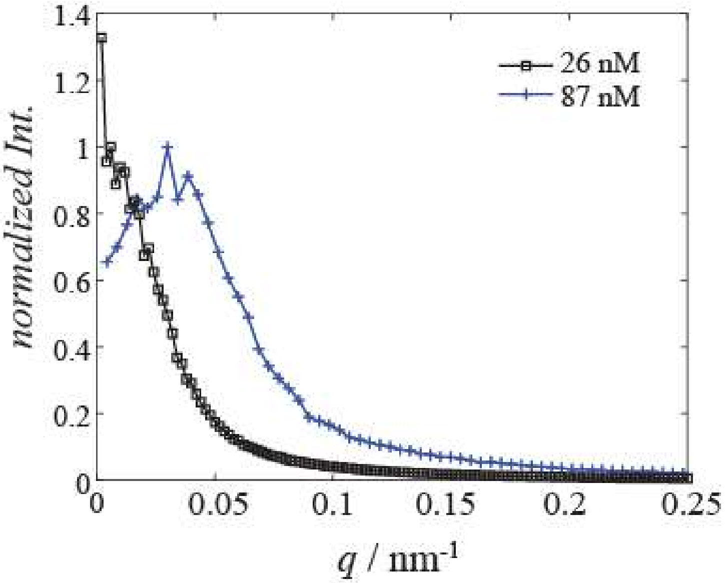


Spectra obtained Radial integration of 2D FFT signals from septins mesh structures on convex regions in SEM images. Radial integration was done by using ImageJ plugin "Radial_Profile_Angle". Integration was done in full angle (360 degrees), and spectra obtained from several images taken at different position in one sample (11 positions for 26 nM and 3 positions for 87 nM) were averaged assuming that the concentration of septins mesh are uniform over the sample. Spectra obtained at different concentration of septins in solution (black square: 26 nM, blue cross: 87 nM) are plotted as a function of $q$ ($nm^{-1}$).

**Supplementary Figure 5**

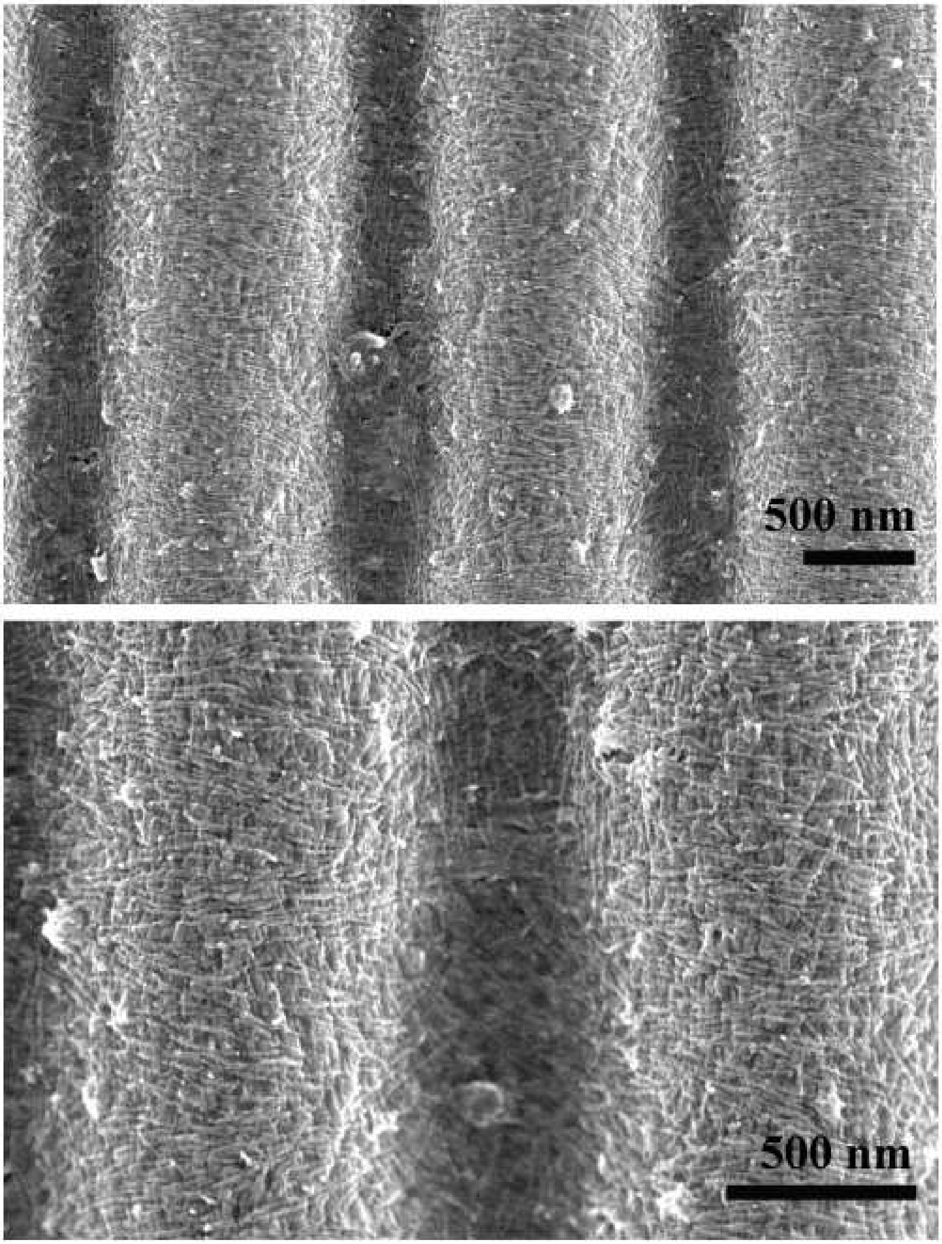


SEM image of septins filaments formed at 44 nM of septins in octamers on lipid bilayer supported by undulated solid substrates with 1.6 μm of periodicity and 0.20 μm of amplitude. Septin's mesh network was observed together with small vesicles bound to the surface.

**Supplementary Figure 6**

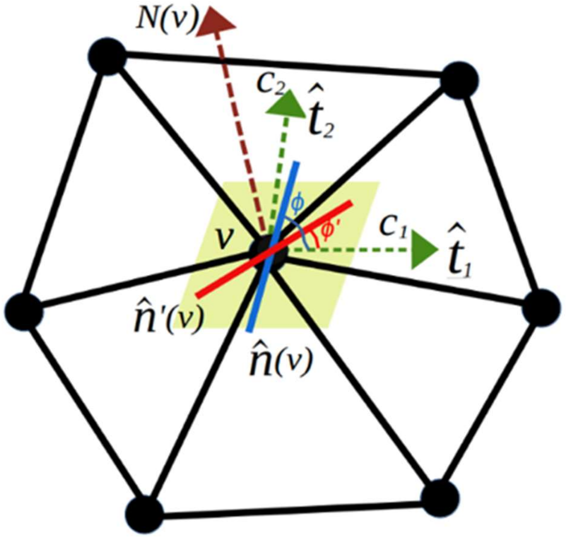


Schematic representation of a small patch of membrane discretized by a network of vertices (solid black circles). At any vertex $v$ the two layers of proteins are modelled by nematic directors $\hat{n}$ and $\hat{n}'$, shown by the blue and red colored rods, respectively, lying on the local tangent plane (in yellow) and characterized by the surface normal N($v$). $c_1$ and $c_2$ are the local principal curvatures at $v$ on the membrane surface along orthogonal tangent vectors $\hat{t}_1$ and $\hat{t}_2$, while $\varphi$ and $\varphi'$ denote the angles between the principal direction $\hat{t}_1$ and the filament orientations $\hat{n}$ and $\hat{n}'$, respectively.

**Supplementary Figure 7**

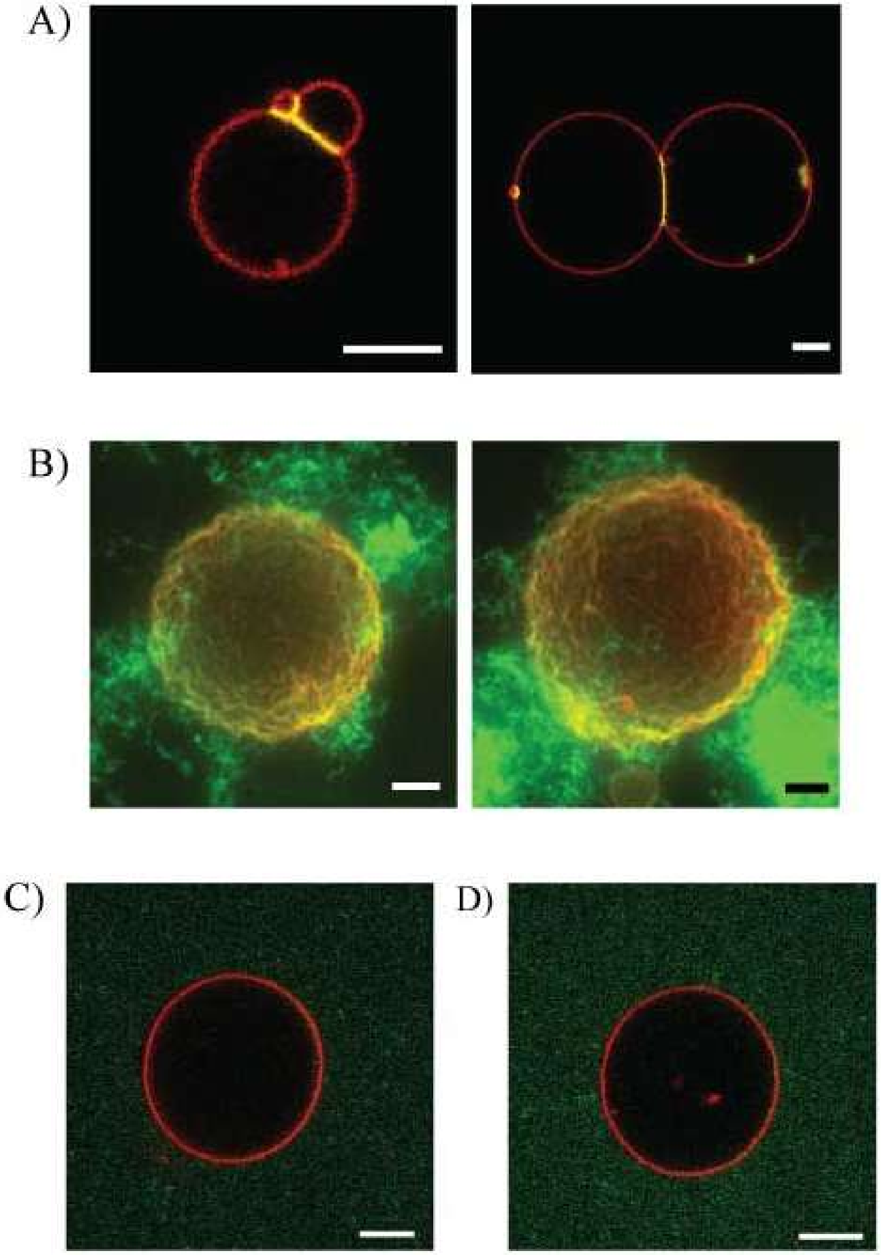


(A) Confocal microscopy images recorded at the equator of GUVs (lipid composition: EggPC 56.5%, Cholesterol 15%, DOPS 10%, DOPE 10%, $PI(4,5)P_2$ 8%, Bodipy TR cermide 0.5%) in the presence of 0.78 nM of human septins octamer-GFP in 70mM NaCl, 10 mM Tris pH 7.8.

(B) 3D reconstruction of confocal microscopy images of GUVs with 520 nM of human septins octamer-GFP in 70mM NaCl, 10 mM Tris pH 7.8.

(C) A confocal microscopy image of a GUV (lipid composition: EggPC 99.5%, Bodipy TR ceramide 0.5%) with 260 nM of human septins octamer in 10 mM Tris pH 7.8 buffer containing 250 mM NaCl.

(D) A confocal microscopy image of a GUV (lipid composition: EggPC 64,5%, Cholesterol 15%, DOPS 10%, DOPE 10%, Bodipy TR cermide 0.5%) with 260 nM of human septins octamer in 10 mM Tris pH 7.8 buffer containing 250 mM NaCl.

Scale bar: 5 μm

**Supplementary Figure 8**

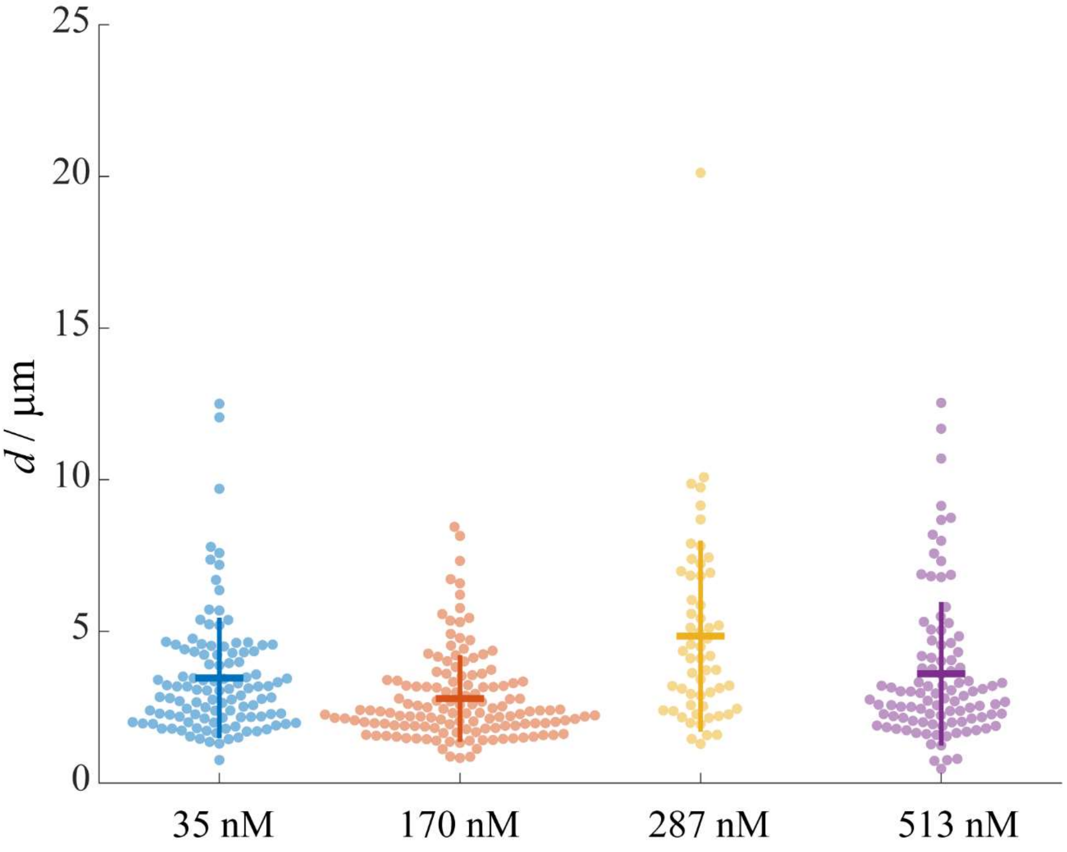


Dependency of distance between peaks of bumps in deformed GUVs on bulk concentration of human septins octamers (nM in octamer). 35 nM of dark human septins (3.5±2.0 μm (mean±SD), 131 counts, $N_{vesicle}$ = 5), 170 nM of human septins-GFP (2.8±1.4 μm (mean±SD), 198 counts, $N_{vesicle}$ = 6), 287 nM of human septins-GFP (4.8±3.2μm (mean±SD), 73 counts, $N_{vesicle}$ = 3), 513 nM of human septins-GFP (3.6±2.4 μm (mean±SD), 202 counts, $N_{vesicle}$ = 4). Perpendicular and horizontal bars show standard deviation (SD) and mean, respectively.

**Supplementary Figure 9**

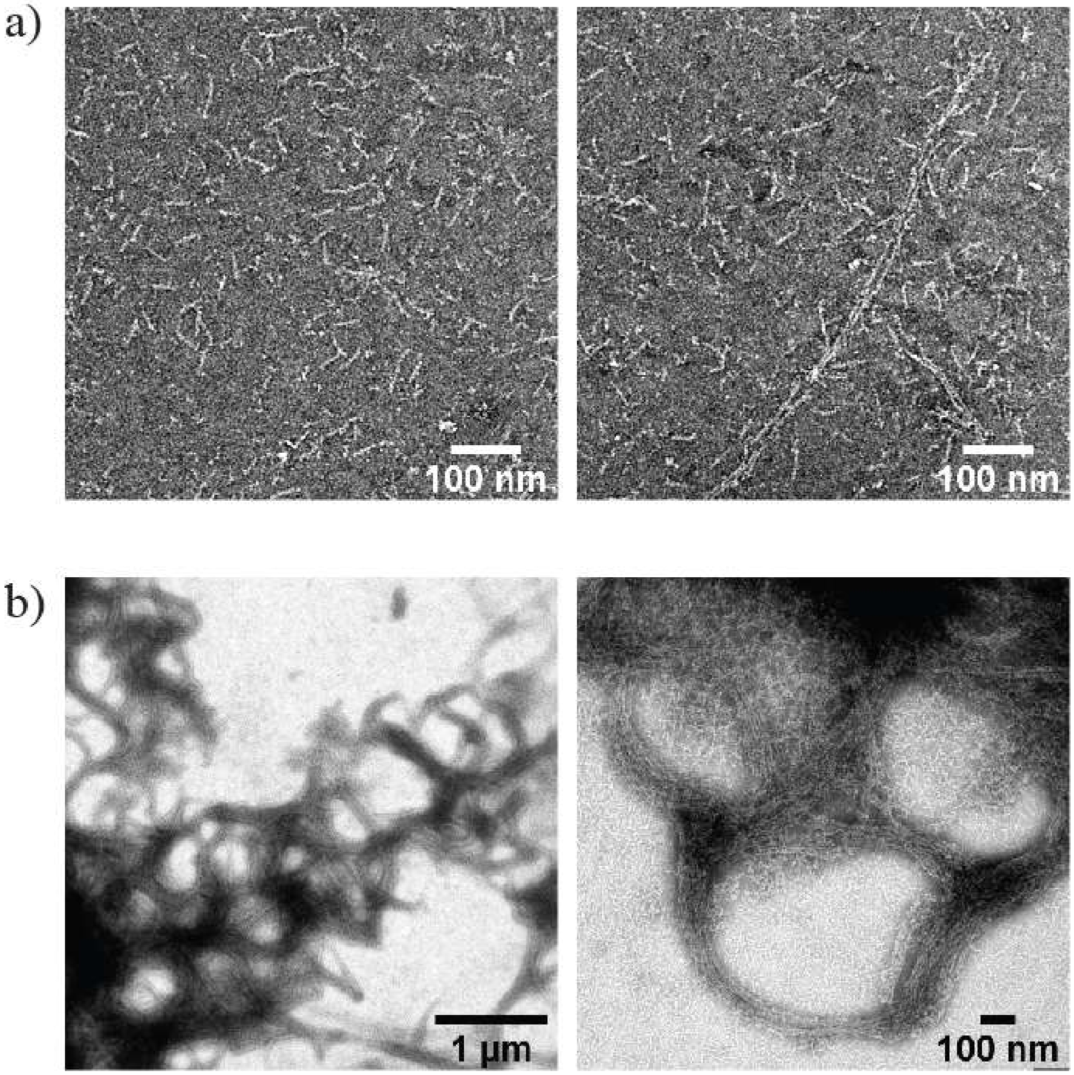


Transmission electron microscopy images of a,b) human septins octamer-GFP in 50 mM Tris pH 8 containing 300 mM KCl, and human septins octamer-GFP after 1h incubation in 10 mM Tris pH 7.8 containing 75 mM NaCl. 4 μL of 0.02 mg/mL for high salt conditions (50 mM Tris pH 8 containing 300 mM KCl) and 0.1 mg/mL for low salt condition (10 mM Tris pH 7.8 containing 75 mM NaCl) were adsorbed for 30s on a carbon-coated electron microscopy grid (Electron Microscopy Sciences, CF300-CU). Grids were then rinsed and negatively stained for 1 min using 1% w/v uranyl formate, and liquids were blotted away with papers and girds were further dried in air. Images were collected using a Tecnai Spirit microscope (Thermo Scientific, FEI) operated at an acceleration voltage of 80kV and equipped with a Quemesa camera (Olympus).

**Supplementary Figure 10**

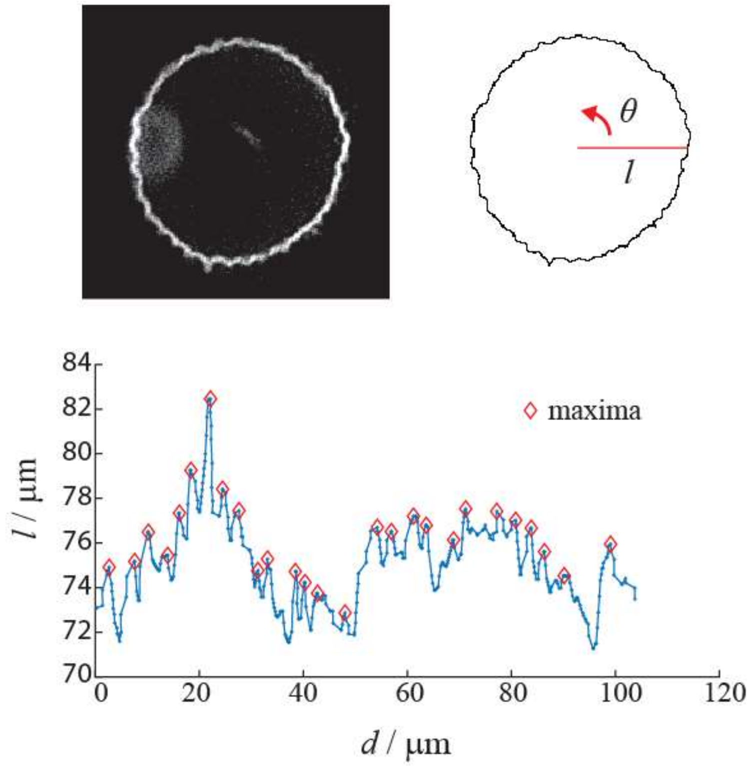


A confocal fluorescence microscopy image of deformed vesicles (top left) and example of image analysis to obtain length *l* (μm) against the angle $\theta$ (top right) according to the x y coordinates of center and perimeter of a vesicle. The angle was converted to the actual distance d by $d = 2\pi R((\theta + 180)/360)\ \times pixel\ size\ (\mu m/pixel)$. Here, *R* is mean radius obtained according to the major and minor length of fitted ellipsoid by (major + minor)/2. Obtained *l* was plotted as a function of *d* (μm) shown in the bottom graph, where obtained local maxima were indicated as red diamonds.

**Supplementary Figure 11**

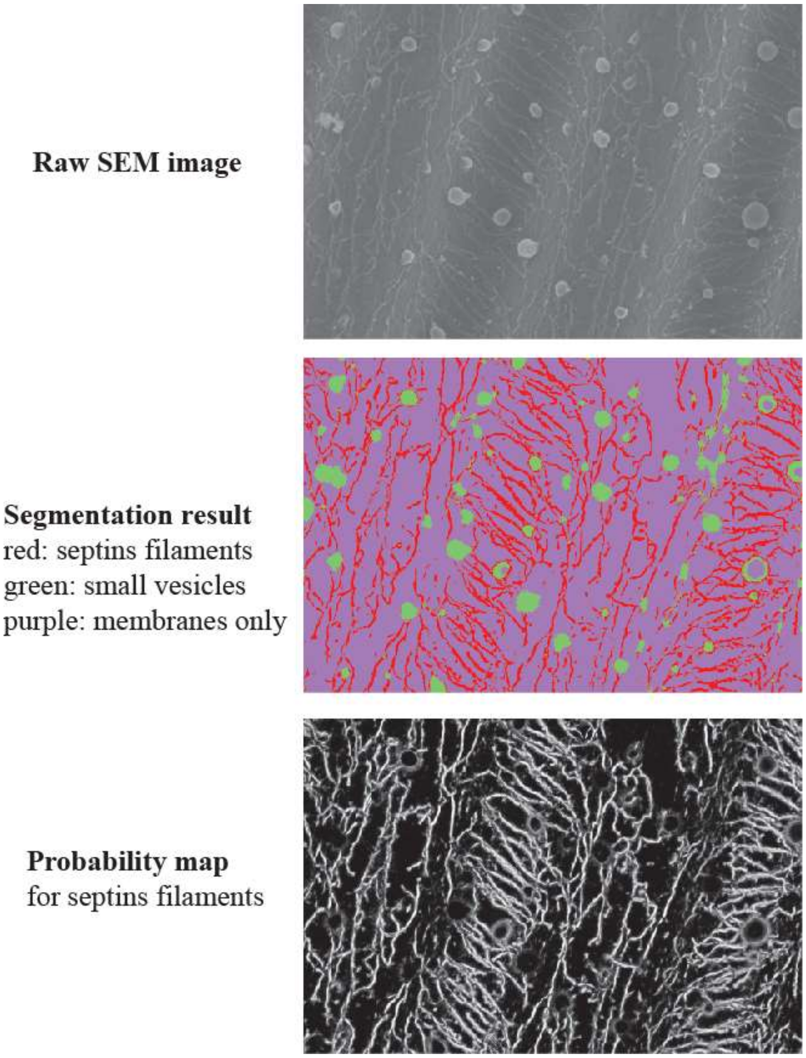


An example of segmentation of a SEM image. Raw image (top) and result of segmentation classified into three categories (septins filaments, small vesicles and membrane only) (middle) and probability map of septins filaments (bottom) by using Weka trainable segmentation in ImageJ software.

**Supplementary Table 1**

| septins conc. | Negative | Positive |
|---|---|---|
| 8.7 nM | $32.2 \pm 2.6$ | $12.4 \pm 2.1$ |
| 26 nM | $45.1 \pm 4.8$ | $28.8 \pm 5.5$ |
| 26 nM (2) | $50.4 \pm 4.6$ | $53.0 \pm 3.4$ |

Surface coverage (%) of undulated supported lipid bilayers by human septins octamer at 8.7 nM and 26 nM of concentration in human septins octamer in solution. Values were obtained by analysis of SEM images where septins filaments were absorbed on lipid bilayer supported by undulated solid substrates with 1.6 μm of periodicity and 0.20 μm of amplitude. SEM images were segmented by using Weka trainable segmentation on imageJ, and probability map of septins filaments were further subjected by thresholding. Threshold image was used to measure the coverage of surface by septins. Measurement was done in 60% of the area near the center of the convex (Positive) or concave (Negative) part, corresponding to positive and negative curvatures, respectively. At 26 nM of septins, we have found roughly two ways of distribution: (1) the place where orientation of filaments is distinctively partitioned, and (2) the place where circular pattern and mesh structures by additional layer of septins were observed. The value corresponds to the mean $\pm$ standard deviation obtained by analysis of several regions (3 regions for 8.7 nM, 6 regions for 26 nM (1) and 13 regions for 26 nM (2)).

**An approximate energy comparison between a smooth surface and valley/hills periodic deformation on fluid membranes at low septin concentrations in simulation**

Here we explain the results found in our simulation, where smooth spherical surface was obtained when 1st and 2nd layer interact with the membrane in equal strength, i.e., $\kappa_{\parallel} = \kappa'_{\parallel}$, while vallay/hill deformations was favorable when 2nd layer weakly interacts, i.e., $\kappa_{\parallel} < \kappa'_{\parallel}$ and $\kappa_{\perp} = \kappa'_{\perp}$. For a sphere (vesicle with radius R) of area A, the constant local curvature $H_{\parallel} = H_{\perp} = \frac{1}{R} = \delta$ is a small number relative to $H_{\parallel} = H_{\perp} = \pm\frac{1}{h}$ for a hill/valley. For a purely convex hill (for example a hemisphere of radius h (with h<<R), the isotropic curvature 1/h > 0 and for a concave valley it is -1/h < 0. We assume that the nematics in the two layers are perpendicular to each other (true at sufficiently high $\epsilon$), and let the intrinsic curvature for septin be $c_{\parallel} = -\alpha$, which is of the same order as $h^{-1}$. While for a sphere, of area A, the Helfrich energy is $8\pi\kappa$, for the undulated vesicle surface of approximately the same total area A, equally distributed between hills and valleys (i.e., ~ A/2 each), the Helfrich energy is higher, namely, $8\pi\kappa\left(\frac{R}{h}\right)^2$. Further, the energy cost from the anisotropic curvature terms, for a sphere, is $E = 2A\left[\frac{\kappa_{\parallel}}{2}(\delta+\alpha)^2 + \frac{\kappa_{\perp}}{2}\delta^2\right] = A[\kappa_{\parallel}(\alpha^2 + 2\alpha\delta) + O(\delta^2)]$. Compared to this, the undulated vesicle surface costs $E = A\left[\frac{\kappa_{\parallel}}{2}(h^{-1}+\alpha)^2 + \frac{\kappa_{\parallel}}{2}(-h^{-1}+\alpha)^2 + \kappa_{\perp}h^{-2}\right] = A[\kappa_{\parallel}(\alpha^2 + h^{-2}) + \kappa_{\perp}h^{-2}]$, which is higher than that of the sphere, since $h^{-1} \gg \delta$. The overall pre-factor 2 in both the expressions is because the 2nd layer, where filament orientations are orthogonal to the 1st layer, gives same contribution as the 1st layer. Note that on a hemisphere $H_{\parallel}$ and $H_{\perp}$do not change with filament orientation. Further, the nematic energy ($\epsilon_{LL}$ term) contribution of the undulated vesicle will be higher than that of the sphere, since surface undulation causes additional topological defects compared to that on a sphere. Note that, since the areas of the spherical vesicle and the deformed vesicle are close their surface tension energies are not significantly different. Besides, septin coated membrane is expected to be stiffer than the bare bilayer membrane (for which κ ~ 10 − 20$k_bT$, we set $\kappa_{\parallel}$=25$k_bT$. We found that setting κ = 0 or κ = 10$k_bT$ in the simulation only makes minor qualitative differences in the shape, in particular, valleys and hills become little flatter with nonzero κ as it opposes higher curvature.
